# COVID-19 Disease Map, a computational knowledge repository of SARS-CoV-2 virus-host interaction mechanisms

**DOI:** 10.1101/2020.10.26.356014

**Authors:** Marek Ostaszewski, Anna Niarakis, Alexander Mazein, Inna Kuperstein, Robert Phair, Aurelio Orta-Resendiz, Vidisha Singh, Sara Sadat Aghamiri, Marcio Luis Acencio, Enrico Glaab, Andreas Ruepp, Gisela Fobo, Corinna Montrone, Barbara Brauner, Goar Frishman, Luis Cristóbal Monraz Gómez, Julia Somers, Matti Hoch, Shailendra Kumar Gupta, Julia Scheel, Hanna Borlinghaus, Tobias Czauderna, Falk Schreiber, Arnau Montagud, Miguel Ponce de Leon, Akira Funahashi, Yusuke Hiki, Noriko Hiroi, Takahiro G. Yamada, Andreas Dräger, Alina Renz, Muhammad Naveez, Zsolt Bocskei, Francesco Messina, Daniela Börnigen, Liam Fergusson, Marta Conti, Marius Rameil, Vanessa Nakonecnij, Jakob Vanhoefer, Leonard Schmiester, Muying Wang, Emily E. Ackerman, Jason Shoemaker, Jeremy Zucker, Kristie Oxford, Jeremy Teuton, Ebru Kocakaya, Gökçe Yağmur Summak, Kristina Hanspers, Martina Kutmon, Susan Coort, Lars Eijssen, Friederike Ehrhart, D. A. B. Rex, Denise Slenter, Marvin Martens, Nhung Pham, Robin Haw, Bijay Jassal, Lisa Matthews, Marija Orlic-Milacic, Andrea Senff Ribeiro, Karen Rothfels, Veronica Shamovsky, Ralf Stephan, Cristoffer Sevilla, Thawfeek Varusai, Jean-Marie Ravel, Rupsha Fraser, Vera Ortseifen, Silvia Marchesi, Piotr Gawron, Ewa Smula, Laurent Heirendt, Venkata Satagopam, Guanming Wu, Anders Riutta, Martin Golebiewski, Stuart Owen, Carole Goble, Xiaoming Hu, Rupert W. Overall, Dieter Maier, Angela Bauch, Benjamin M. Gyori, John A. Bachman, Carlos Vega, Valentin Grouès, Miguel Vazquez, Pablo Porras, Luana Licata, Marta Iannuccelli, Francesca Sacco, Anastasia Nesterova, Anton Yuryev, Anita de Waard, Denes Turei, Augustin Luna, Ozgun Babur, Sylvain Soliman, Alberto Valdeolivas, Marina Esteban- Medina, Maria Peña-Chilet, Kinza Rian, Tomáš Helikar, Bhanwar Lal Puniya, Dezso Modos, Agatha Treveil, Marton Olbei, Bertrand De Meulder, Aurélien Dugourd, Aurélien Naldi, Vincent Noë, Laurence Calzone, Chris Sander, Emek Demir, Tamas Korcsmaros, Tom C. Freeman, Franck Augé, Jacques S. Beckmann, Jan Hasenauer, Olaf Wolkenhauer, Egon L. Wilighagen, Alexander R. Pico, Chris T. Evelo, Marc E. Gillespie, Lincoln D. Stein, Henning Hermjakob, Peter D’Eustachio, Julio Saez-Rodriguez, Joaquin Dopazo, Alfonso Valencia, Hiroaki Kitano, Emmanuel Barillot, Charles Auffray, Rudi Balling, Reinhard Schneider, the COVID-19 Disease Map Community

**Affiliations:** Luxembourg Centre for Systems Biomedicine, University of Luxembourg, Esch-sur-Alzette, Luxembourg; Department of Biology, Univ. Evry, University of Paris-Saclay, GenHotel, Genopole, 91025, Evry, France; Lifeware Group, Inria Saclay-Ile de France, Palaiseau 91120, France; Institut Curie, PSL Research University, Paris, France; INSERM, U900, Paris, France; MINES ParisTech, PSL Research University, Paris, France; Integrative Bioinformatics, Inc., 346 Paul Ave, Mountain View, CA, USA; Institut Pasteur, HIV, Inflammation and Persistence Unit, Paris, France; Bio Sorbonne Paris Cité, Université de Paris, Paris, France; Inserm- Institut national de la santé et de la recherche médicale. Saint-Louis Hospital 1 avenue Claude Vellefaux Pavillon Bazin 75475 Paris; Institute of Experimental Genetics (IEG), Helmholtz Zentrum München-German Research Center for Environmental Health (GmbH), Ingolstädter Landstraße 1, D-85764 Neuherberg, Germany; Oregon Health & Sciences University; Department of Molecular and Medical Genetics; 3222 SW Research Drive, Portland, Oregon, U.S.A 97239; Department of Systems Biology and Bioinformatics, University of Rostock, 18051 Rostock, Germany; Department of Computer and Information Science, University of Konstanz, Konstanz, Germany; Monash University, Faculty of Information Technology, Department of Human-Centred Computing, Wellington Rd, Clayton VIC 3800, Australia; Barcelona Supercomputing Center (BSC), Barcelona, Spain; Keio University, Department of Biosciences and Informatics, 3-14-1 Hiyoshi Kouhoku-ku Yokohama Japan 223-8522; Sanyo-Onoda City University, Faculty of Pharmaceutical Sciences, University St.1-1-1, Yamaguchi, Japan 756-0884; Computational Systems Biology of Infections and Antimicrobial-Resistant Pathogens, Institute for Bioinformatics and Medical Informatics (IBMI), University of Tübingen, 72076 Tübingen, Germany; Department of Computer Science, University of Tübingen, 72076 Tübingen, Germany; German Center for Infection Research (DZIF), partner site Tübingen, Germany; Riga Technical University, Institute of Applied Computer Systems,1 Kalku Street, LV-1658 Riga, Latvia; Sanofi R&D, Translational Sciences, 1 av Pierre Brossolette 91395 Chilly-Mazarin France; Dipartimento di Epidemiologia Ricerca Pre-Clinica e Diagnostica Avanzata, National Institute for Infectious Diseases ’Lazzaro Spallanzani’ I.R.C.C.S., Rome, Italy; COVID 19 INMI Network Medicine for IDs Study Group, National Institute for Infectious Diseases ’Lazzaro Spallanzani’ I.R.C.C.S., Rome, Italy; Bioinformatics Core Facility, Universitätsklinikum Hamburg-Eppendorf, Martinistraße 52, 20246 Hamburg, Germany; The University of Edinburgh, Royal (Dick) School of Veterinary Medicine, Easter Bush Campus, Midlothian, EH25 9RG; Faculty of Mathematics and Natural Sciences, University of Bonn, Bonn, Germany; Helmholtz Zentrum München - German Research Center for Environmental Health, Institute of Computational Biology, 85764 Neuherberg, Germany; Technische Universität München, Center for Mathematics, Chair of Mathematical Modeling of Biological Systems, 85748 Garching, Germany; Department of Chemical and Petroleum Engineering, University of Pittsburgh; Department of Computational and Systems Biology, University of Pittsburgh; Pacific Northwest National Laboratory, 902 Battelle Boulevard, Richland, WA, US; Ankara University, Stem Cell Institute, Ceyhun Atif Kansu St. No: 169 06520 Cevizlidere, Ankara, Turkey; Institute of Data Science and Biotechnology, Gladstone Institutes, San Francisco, CA 94158, US; Department of Bioinformatics - BiGCaT, NUTRIM, Maastricht University, Maastricht, The Netherlands; Maastricht Centre for Systems Biology (MaCSBio), Maastricht University, Maastricht, The Netherlands; Maastricht University Medical Centre, Universiteitssingel 50, 6229 ER Maastricht, The Netherlands; Center for Systems Biology and Molecular Medicine, Yenepoya (Deemed to be University), Mangalore 575018, India; Ontario Institute for Cancer Research, MaRS Centre, 661 University Avenue, Suite 510, Toronto, Ontario, Canada M5G 0A3; NYU Grossman School of Medicine, New York NY 10016, US; Universidade Federal do Paraná, Brasil; EMBL-EBI, Molecular Systems, Wellcome Genome Campus, Hinxton, Cambridgeshire, CB10 1SD; University of Lorraine, INSERM UMR_S 1256, Nutrition, Genetics, and Environmental Risk Exposure (NGERE), Faculty of Medicine of Nancy, F-54000 Nancy, France; Laboratoire de génétique médicale, CHRU Nancy, Nancy, France; The University of Edinburgh, Queen’s Medical Research Institute. 47 Little France Crescent, Edinburgh, EH16 4TJ; Senior Research Group in Genome Research of Industrial Microorganisms, Center for Biotechnology, Bielefeld University, Universitätsstraße 27, 33615 Bielefeld, Germany; Department of Surgical Science, Uppsala University, Sweden; Poznan University of Technology, Institute of Computing Science, ul. Piotrowo 2, 60-965 Poznan, Poland; Department of Medical Informatics and Clinical Epidemiology, Oregon Health & Science University, 3181 S.W. Sam Jackson Park Road, Portland, OR 97239-3098, USA; Heidelberg Institute for Theoretical Studies (HITS), Schloss-Wolfsbrunnenweg 35, D-69118 Heidelberg, Germany; The University of Manchester, Department of Computer Science, Oxford Road, Manchester, M13 9PL, UK; German Center for Neurodegenerative Diseases (DZNE) Dresden, Tatzberg 41, 01307 Dresden, Germany; Center for Regenerative Therapies Dresden (CRTD), Technische Universität Dresden, Fetscherstraße 105, 01307 Dresden, Germany; Biomax Informatics AG, Robert-Koch-Str. 2, 82152 Planegg, Germany; Harvard Medical School, Laboratory of Systems Pharmacology, 200 Longwood Avenue, Boston, MA, US; Open Targets, Wellcome Genome Campus, Hinxton, Cambridgeshire, CB10 1SD; University of Rome Tor Vergata, Department of Biology, Via della Ricerca Scientifica 1, 00133 Rome, Italy; Elsevier, 1600 John F Kennedy Blvd #1800, Philadelphia, PA 19103, US; Elsevier, Research Collaborations Unit, 71 Hanley Lane, Jericho, VT 05465, US; Heidelberg University, Institute for Computational Biomedicine, BQ 0053, Im Neuenheimer Feld 267, 69120 Heidelberg, Germany; cBio Center, Divisions of Biostatistics and Computational Biology, Department of Data Sciences, Dana-Farber Cancer Institute, Boston, MA, 02215, US; Department of Cell Biology, Harvard Medical School, Boston, MA, 02115, US; University of Massachusetts Boston, Computer Science Department, 100 William T, Morrissey Blvd, Boston, MA 02125, US; Clinical Bioinformatics Area, Fundación Progreso y Salud (FPS), Hospital Virgen del Rocio, Sevilla, 41013, Spain; Computational Systems Medicine group, Institute of Biomedicine of Seville (IBIS). Hospital Virgen del Rocio, Sevilla 41013, Spain; Bioinformatics in Rare Diseases (BiER). Centro de Investigación Biomédica en Red de Enfermedades Raras (CIBERER), FPS, Hospital Virgen del Rocío, 41013, Sevilla, Spain; University of Nebraska-Lincoln, Department of Biochemistry, 1901 Vine St., Lincoln, NE, 68588, US; Quadram Institute Bioscience, Rosalind Franklin Road, Norwich Research Park, Norwich, NR4 7UQ, UK; Earlham Institute, Norwich Research Park, Norwich, NR4 7UZ, UK; European Institute for Systems Biology and Medicine (EISBM), Vourles, France; Institute of Experimental Medicine and Systems Biology, Faculty of Medicine, RWTH, Aachen University, Aachen, Germany; The Roslin Institute, University of Edinburgh EH25 9RG; University of Lausanne, Lausanne, Switzerland; Interdisciplinary Research Unit Mathematics and Life Sciences, University of Bonn, Germany; StJohn’s University College of Pharmacy and Health Sciences, Queens, NY USA; FPS/ELIXIR-es, Hospital Virgen del Rocío, Sevilla, 42013, Spain; Institució Catalana de Recerca i Estudis Avançats (ICREA), Barcelona, Spain; Systems Biology Institute, Tokyo Japan; Okinawa Institute of Science and Technology Graduate School

## Abstract

We describe a large-scale community effort to build an open-access, interoperable, and computable repository of COVID-19 molecular mechanisms - the COVID-19 Disease Map. We discuss the tools, platforms, and guidelines necessary for the distributed development of its contents by a multi-faceted community of biocurators, domain experts, bioinformaticians, and computational biologists. We highlight the role of relevant databases and text mining approaches in enrichment and validation of the curated mechanisms. We describe the contents of the Map and their relevance to the molecular pathophysiology of COVID-19 and the analytical and computational modelling approaches that can be applied for mechanistic data interpretation and predictions. We conclude by demonstrating concrete applications of our work through several use cases and highlight new testable hypotheses.

## 1. Introduction

The coronavirus disease 2019 (COVID-19) pandemic due to severe acute respiratory syndrome coronavirus 2 (SARS-CoV-2) already resulted in the infection of over 106 million people worldwide, of whom 2.3 million have died^1^. The molecular pathophysiology that links SARS-CoV-2 infection to the clinical manifestations and course of COVID-19 is complex and spans multiple biological pathways, cell types and organs [1,2]. To gain insight into this network of molecular mechanisms we need knowledge collected from the scientific literature and bioinformatic databases, integrated using formal systems biology standards. A repository of such computable knowledge will support data analysis and predictive modelling.

With this goal in mind, we initiated a collaborative effort involving over 230 biocurators, domain experts, modellers and data analysts from 120 institutions in 30 countries to develop the COVID-19 Disease Map, an open-access collection of curated computational diagrams and models of molecular mechanisms implicated in the disease [3].

To this end, we aligned the biocuration efforts of the Disease Maps Community [4,5], Reactome [6], and WikiPathways [7] and developed common guidelines utilising standardised encoding and annotation schemes, based on community-developed systems biology standards [8–10], and persistent identifier repositories [11]. Moreover, we integrated relevant knowledge from public repositories [12–15] and text mining resources, providing a means to update and refine the contents of the Map. The fruit of these efforts was a series of pathway diagrams describing key events in the COVID-19 infectious cycle and host response.

This comprehensive diagrammatic description of disease mechanisms is machine-readable and computable. This allows us to develop novel bioinformatics workflows, creating executable networks for analysis and prediction. In this way, the Map is both human and machine-readable, lowering communication barriers between biocurators, domain experts, and computational biologists significantly. Computational modelling, data analysis, and their informed interpretation using the contents of the Map have the potential to identify molecular signatures of disease predisposition and development, and to suggest drug repositioning for improving current treatments.

The current COVID-19 Disease Map is a collection of 41 diagrams containing 1836 interactions between 5499 elements, supported by 617 publications and preprints. The summary of diagrams available in the COVID-19 Disease Map can be found online^2^in Supplementary Material 1. The Map is a constantly evolving resource, refined and updated by ongoing efforts of biocuration, sharing and analysis. Here, we report its current status.

In Section 2, we explain the set up of our community effort to construct the interoperable content of the resource, involving biocurators, domain experts and data analysts. In Section 3, we demonstrate that the scope of the biological maps in the resource reflects the state-of-the-art about the molecular biology of COVID-19. Next, we outline analytical workflows that can be used on the contents of the Map, including initial, preliminary outcomes of two such workflows, discussed in detail as use cases in Section 4. We conclude in Section 5 with an outlook to further development of the COVID-19 map and the utility of the entire resource in future efforts to build and apply disease-relevant computational repositories.

## 2. Building and sharing the interoperable content

The COVID-19 Disease Map project involves: (i) biocurators, (ii) domain experts, and (iii) analysts and modellers:

i. Biocurators develop a collection of systems biology diagrams focused on the molecular mechanisms of SARS-CoV-2.
ii. Domain experts refine the contents of the diagrams, supported by interactive visualisation and annotations.
iii. Analysts and modellers develop computational workflows to generate hypotheses and predictions about the mechanisms encoded in the diagrams.

All three groups have an essential role in the process of building the Map, by providing content, refining it, and defining its computational use. Figure 1 illustrates the ecosystem of the COVID-19 Disease Map Community, highlighting the roles of different participants, available format conversions, interoperable tools, and downstream uses. Information about the community members and their contributions is disseminated via the FAIRDOMHub [16], so that content distributed across different collections can be uniformly referenced.

**Figure 1:**
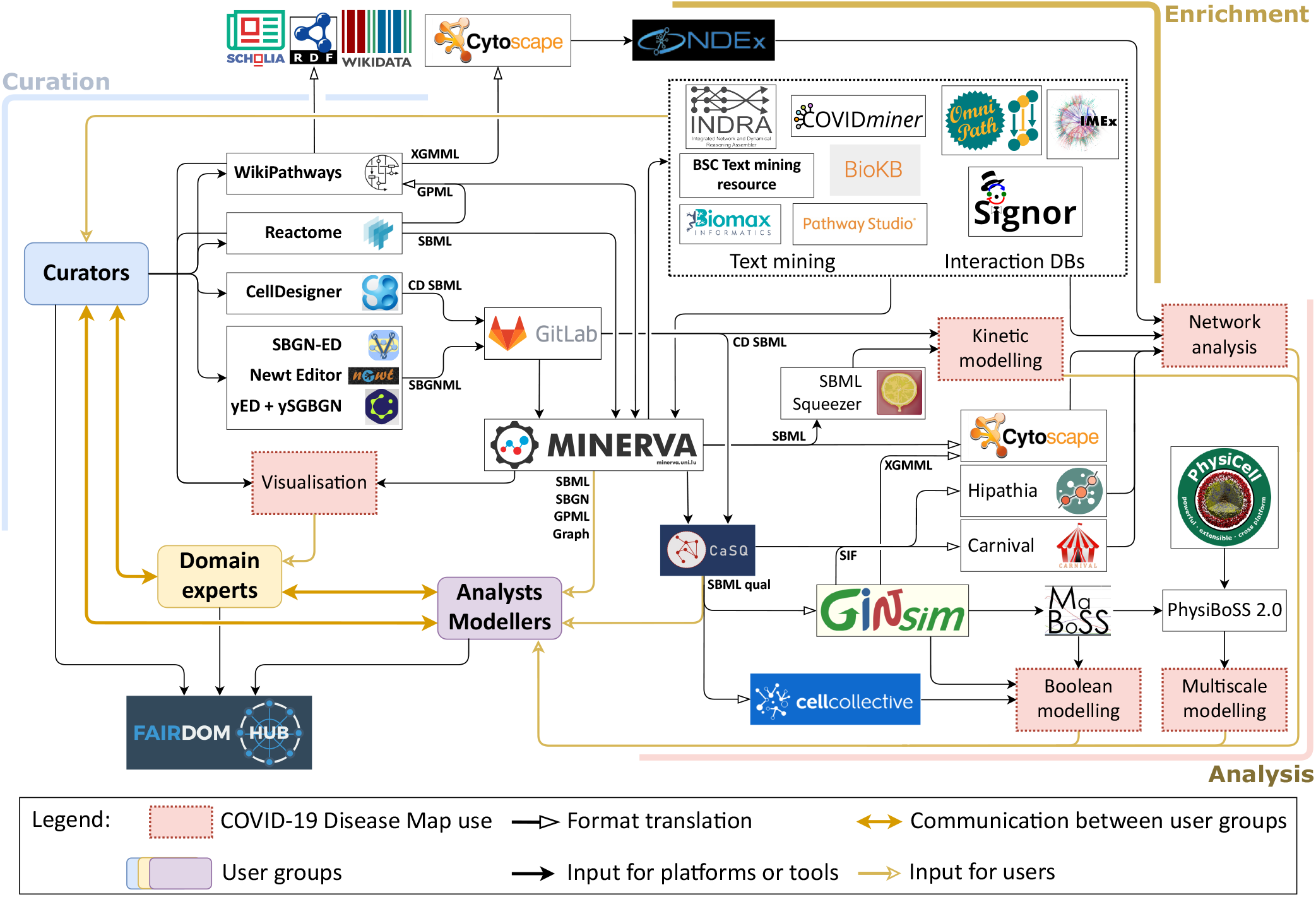
The ecosystem of the COVID-19 Disease Map Community. The main groups of COVID-19 Disease Map Community are biocurators, domain experts, analysts, and modellers; communicating to refine, interpret and apply COVID-19 Disease Map diagrams. These diagrams are created and maintained by biocurators, following pathway database workflows or standalone diagram editors, and reviewed by domain experts. The content is shared via pathway databases or a GitLab repository; all can be enriched by integrated resources of text mining and interaction databases. The COVID-19 Disease Map diagrams are available in several layout-aware systems biology formats and integrated with external repositories, allowing a range of computational analyses, including network analysis and Boolean, kinetic or multiscale simulations.

### 2.1 Creating and accessing the diagrams

The biocurators of the COVID-19 Disease Map diagrams follow the guidelines developed by the Community, and specific workflows of WikiPathways [7] and Reactome [6]. The biocurators build literature-based systems biology diagrams, representing the molecular processes implicated in COVID-19 pathophysiology, their complex regulation and the phenotypic outcomes. These diagrams are the main building blocks of the Map, composed of biochemical reactions and interactions (altogether called interactions) taking place between different types of molecular entities in various cellular compartments. As multiple teams work on related topics, biocurators can provide an expert review across pathways and across platforms. This is possible, as all platforms offer intuitive visualisation, interpretation, and analysis of pathway knowledge to support basic and clinical research, genome analysis, modelling, systems biology, and education. Table 1 lists information about the created content. For more details see Supplementary Material 1.

**Table 1.**
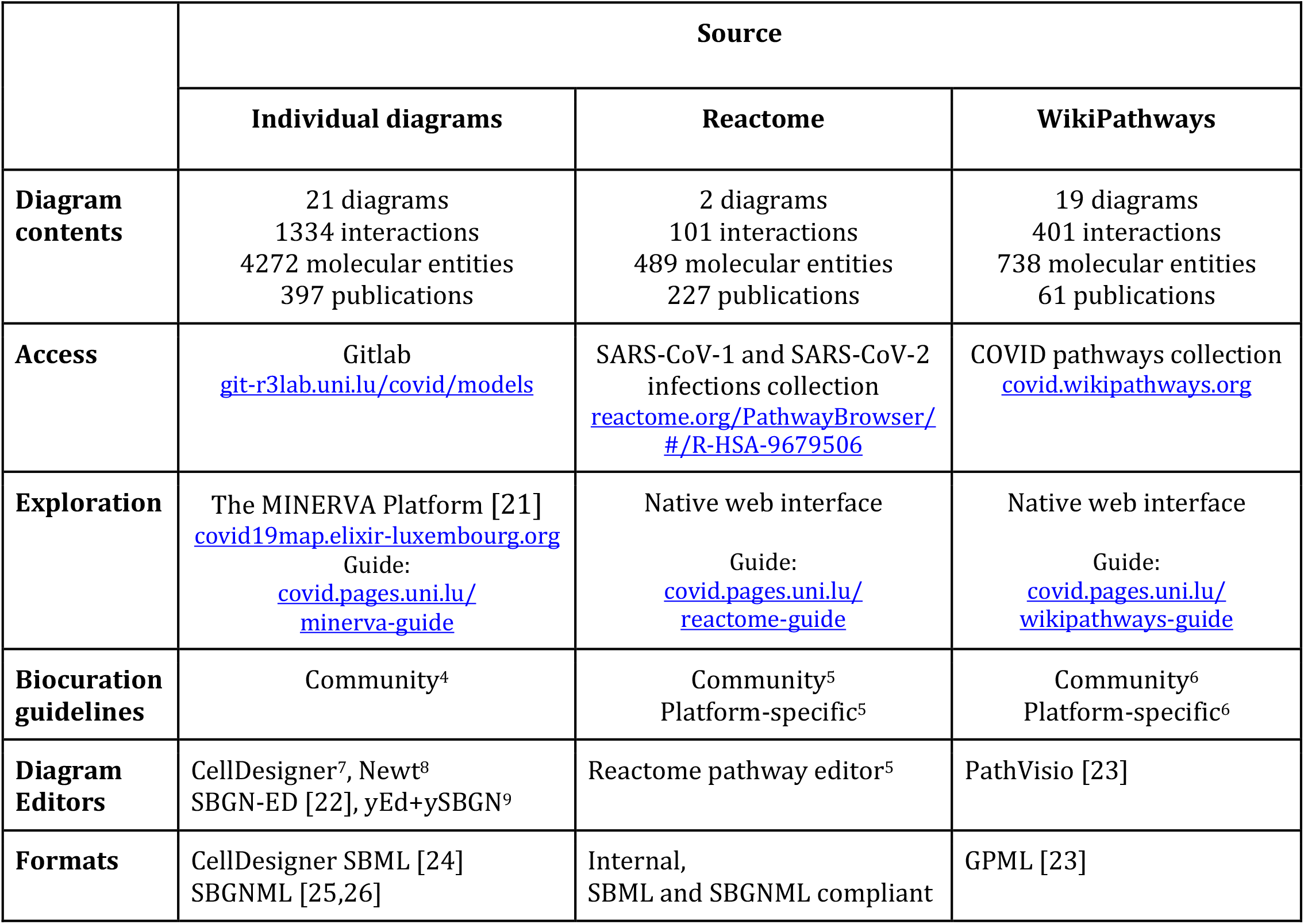
COVID-19 Disease Map contents. The table summarises biocuration resources and content of the Map across three main parts of the repository. All diagrams are listed in Supplementary Table 1. Available online at https://covid.pages.uni.lu/map_contents.

|  | Source |  |  |
| --- | --- | --- | --- |
|  | Individual diagrams | Reactome | WikiPathways |
| **Diagram contents** | 21 diagrams<br>1334 interactions<br>4272 molecular entities<br>397 publications | 2 diagrams<br>101 interactions<br>489 molecular entities<br>227 publications | 19 diagrams<br>401 interactions<br>738 molecular entities<br>61 publications |
| **Access** | Gitlab<br>git-r3lab.uni.lu/covid/models | SARS-CoV-1 and SARS-CoV-2 infections collection<br>reactome.org/PathwayBrowser/#/R-HSA-9679506 | COVID pathways collection<br>covid.wikipathways.org |
| **Exploration** | The MINERVA Platform [21]<br>covid19map.elixir-luxembourg.org<br>Guide:<br>covid.pages.uni.lu/minerva-guide | Native web interface<br>Guide:<br>covid.pages.uni.lu/reactome-guide | Native web interface<br>Guide:<br>covid.pages.uni.lu/wikipathways-guide |
| **Biocuration guidelines** | Community<sup>4</sup> | Community<sup>5</sup><br>Platform-specific<sup>5</sup> | Community<sup>6</sup><br>Platform-specific<sup>6</sup> |
| **Diagram Editors** | CellDesigner<sup>7</sup>, Newt<sup>8</sup><br>SBGN-ED [22], yEd+ySBGN<sup>9</sup> | Reactome pathway editor<sup>5</sup> | PathVisio [23] |
| **Formats** | CellDesigner SBML [24]<br>SBGNML [25,26] | Internal,<br>SBML and SBGNML compliant | GPML [23] |

Both interactions and interacting entities are annotated following a uniform, persistent identification scheme, using either MIRIAM or Identifiers.org [17], and the guidelines for annotations of computational models [18]. Viral protein interactions are explicitly annotated with their taxonomy identifiers to highlight findings from strains other than SARS-CoV-2. Moreover, tools like ModelPolisher [19], SBMLsqueezer [20] or MEMOTE^3^help to automatically complement the annotations in the SBML format and validate the model (see also Supplementary Material 2).

### 2.2 Enrichment using knowledge from databases and text mining

The knowledge on COVID-19 mechanisms is rapidly evolving, as demonstrated by the rapid growth of the COVID-19 Open Research Dataset (CORD-19) dataset, a source of scientific manuscripts and metadata on COVID-19 and related coronavirus research [27]. CORD-19 currently contains over 130,000 articles and preprints, over four times more than when it was introduced^10^. In such a quickly evolving environment, biocuration efforts need to be supported by repositories of structured knowledge about molecular mechanisms relevant for COVID-19, like molecular interaction databases, or text mining resources. Contents of such repositories may suggest improvements in the existing COVID-19 Disease Map diagrams, or establish a starting point for developing new pathways (see Section “Biocuration of database and text mining content”).

#### Interaction and pathway databases

Interaction and pathway databases contain structured and annotated information on protein interactions or causal relationships, while interaction databases focus on pairs of molecules, offering broad coverage of literature-reported findings, pathway databases describe biochemical processes and their regulations, supported by diagrams. Both types of resources are valuable inputs for COVID-19 Disease Map biocurators, given the comparability of identifiers used for molecular annotations, and the reference to publications used for defining an interaction or building a pathway. Table 2 summarises open-access resources supporting the biocuration of the Map. See Supplementary Materials [tools] for their detailed description.

**Table 2.**
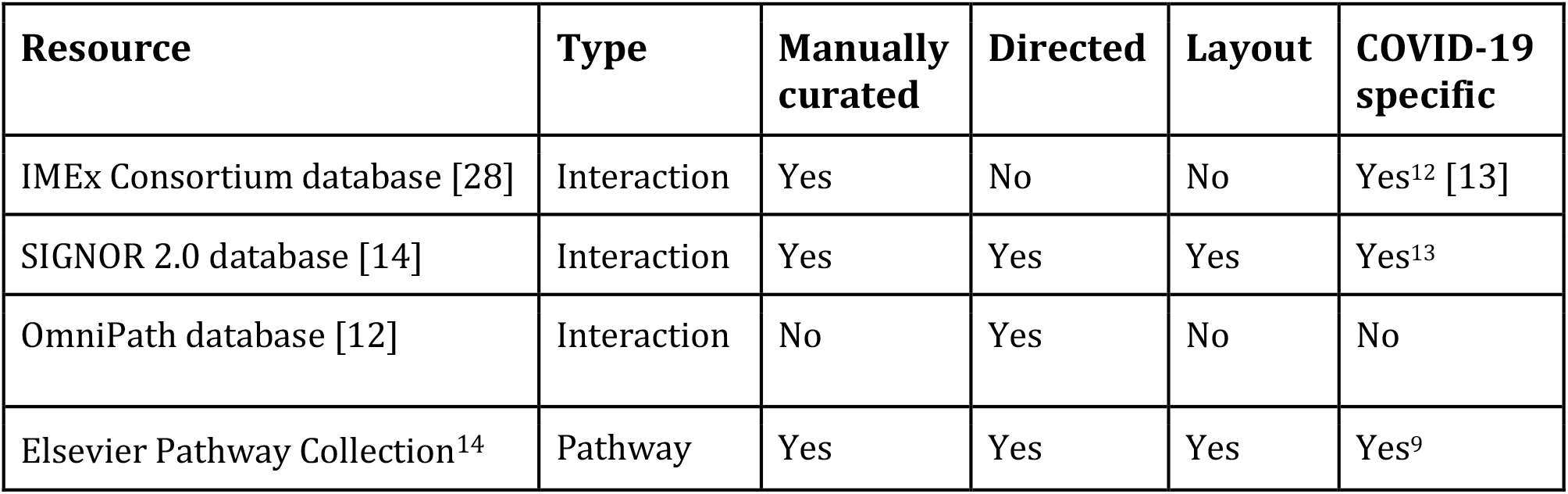
Resources supporting biocuration of the COVID-19 Disease Map. They include (i) collections of COVID-19 interactions, published by the IMEx Consortium [13] and SIGNOR 2.0 [14], (ii) a non-COVID interaction database OmniPath [12] and (iii) the Elsevier Pathway Collection, a manually reconstructed open-access dataset of annotated pathway diagrams for COVID-19^11^.

| Resource | Type | Manually curated | Directed | Layout | COVID-19 specific |
| --- | --- | --- | --- | --- | --- |
| IMEx Consortium database [28] | Interaction | Yes | No | No | Yes<sup>12</sup> [13] |
| SIGNOR 2.0 database [14] | Interaction | Yes | Yes | Yes | Yes<sup>13</sup> |
| OmniPath database [12] | Interaction | No | Yes | No | No |
| Elsevier Pathway Collection<sup>14</sup> | Pathway | Yes | Yes | Yes | Yes<sup>9</sup> |

<sup>10</sup> https://www.semanticscholar.org/cord19/download (accessed on 20.10.2020)
<sup>11</sup> https://data.mendeley.com/datasets/h9vs5s8fz2/draft?a=f40961bb-9798-4fd1-8025-e2a3ba47b02e
<sup>12</sup> [https://www.ebi.ac.uk/intact/imex/main.xhtml?query=annot:"dataset:coronavirus"](https://www.ebi.ac.uk/intact/imex/main.xhtml?query=annot:)
<sup>13</sup> https://signor.uniroma2.it/covid/
<sup>14</sup> https://pathwaystudio.com

#### Text mining resources

Text-mining approaches can help to sieve through such rapidly expanding literature with natural language processing (NLP) algorithms based on semantic modelling, ontologies, and linguistic analysis to automatically extract and annotate relevant sentences, biomolecules, and their interactions. This scope was recently extended to pathway figure mining, decoding pathway figures into their computable representations [29]. Altogether, these automated workflows lead to the construction of knowledge graphs: semantic networks incorporating ontology concepts, unique biomolecule references, and their interactions extracted from abstracts or full-text documents [30].

The COVID-19 Disease Map Project integrates open-access text mining resources, INDRA [31], BioKB^15^, AILANI COVID-19^16^, and PathwayStudio^14^. All platforms offer keyword-based search allowing interactive exploration. Additionally, the Map benefits from an extensive protein-protein interaction network (PPI)^17^generated with a custom text-mining pipeline using OpenNLP^18^and GNormPlus [32]. This pipeline was applied to the CORD-19 dataset and the collection of MEDLINE abstracts associated with the genes in the SARS-CoV-2 PPI network [33] using the Entrez Gene Reference-Into-Function (GeneRIF). For detailed descriptions of the resources, see Supplementary Material 3.

#### Biocuration using database and text mining content

Molecular interactions from databases and knowledge graphs from text mining resources discussed above (from now on called altogether ‘knowledge graphs’) have a broad coverage at the cost of depth of mechanistic representation. This content can be used by the biocurators to build and update the systems biology focused diagrams. Biocurators can use this content in three main ways: by visual exploration, by programmatic comparison, and by direct incorporation of the content.

First, the biocurators can visually explore the contents of the knowledge graphs using available search interfaces to locate new knowledge and encode it in the diagrams. Moreover, solutions like COVIDminer project^19^, PathwayStudio^14^ and AILANI offer a visual representation of a group of interactions for a better understanding of their biological context, allowing search by interactions, rather than just isolated keywords. Finally, INDRA and AILANI offer assistant bots that respond to natural language queries and return meaningful answers extracted from knowledge graphs.

Second, programmatic access and reproducible exploration of the knowledge graphs is possible via data endpoints: SPARQL for BioKB and Application Programming Interfaces for INDRA, AILANI, and Pathway Studio. Users can programmatically submit keyword queries and retrieve functions, interactions, pathways, or drugs associated with submitted gene lists. This way, otherwise time-consuming tasks like an assessment of completeness of a given diagram, or search for new literature evidence, can be automated to a large extent.

Finally, biocurators can directly incorporate the content of knowledge graphs into SBML format using BioKC [34]. Additionally, the contents of the Elsevier COVID-19 Pathway Collection can be translated to SBGNML^20^ preserving the layout of the diagrams. The SBGNML content can then be converted into other diagram formats used by biocurators (see Section 2.3 below).

### 2.3 Interoperability of the diagrams and annotations

The biocuration of the COVID-19 Disease Map is distributed across multiple teams, using varying tools and associated systems biology representations. This requires a common approach to annotations of evidence, biochemical reactions, molecular entities and their interactions. Moreover, interoperability of layout-aware formats is needed for comparison and integration of the diagrams in the Map.

#### Layout-aware formats for molecular mechanisms

The COVID-19 Disease Map diagrams are encoded in one of three layout-aware formats for standardised representation of molecular interactions: SBML^21^ [35–37], SBGNML [26], and GPML [23]. These XML-based formats focus to a varying degree on user-friendly graphical representation, standardised visualisation, and support of computational workflows. For the detailed description of the formats, see Supplementary Material 1.

Each of these three languages has a different focus: SBML emphasises standardised representation of the data model underlying molecular interactions, SBGNML provides a standardised graphical representation of molecular processes, while GPML allows for a partially standardised representation of uncertain biological knowledge. Nevertheless, all three formats are centred around molecular interactions, provide a constrained vocabulary to encode element and interaction types, encode layout of their diagrams and support stable identifiers for diagram components. These shared properties, supported by a common ontology^22^ [38], allow cross-format mapping and enable translation of key properties between the formats. Therefore, when developing the contents of the Map, biocurators use the tools they are familiar with, facilitating this distributed task.

#### Format interoperability

The COVID-19 Disease Map Community ecosystem of tools and resources (see Figure 1) ensures interoperability between the three layout-aware formats for molecular mechanisms: SBML, SBGNML, and GPML. Essential elements of this setup are tools capable of providing cross-format translation functionality [39,40] and supporting harmonised visualisation processing. Another essential translation interface is a representation of Reactome pathways in WikiPathways GPML [41] and SBML. The SBML export of Reactome content has been optimised in the context of this project and facilitates integration with the other COVID-19 Disease Map software components.

The contents of the COVID-19 Disease Map diagrams can be directly transformed into inputs of computational pipelines and data repositories. Besides the direct use of SBML format in kinetic simulations, CellDesigner SBML files can be transformed into SBML qual [42] using CaSQ [43], enabling Boolean modelling-based simulations (see also Supplementary Material 3). CaSQ preserves annotations and layout information for transparency and reusability of the models. In parallel, CaSQ converts the diagrams to the SIF format^23^, supporting pathway modelling workflows using simplified interaction networks. Notably, the GitLab repository features an automated translation of stable versions of diagrams into SBML qual. Finally, translation of the diagrams into XGMML format (the eXtensible Graph Markup and Modelling Language) using Cytoscape [44] or GINSim [45] allows for network analysis and interoperability with molecular interaction repositories [46].

## 3. Structure and scope of the COVID-19 Disease Map

The COVID-19 Disease Map is the product of a large-scale community effort. It was built bottom-up, exploiting a rich bioinformatics framework, on a skeleton provided from previous extensive studies of other coronaviruses [47] and contextualised with data emerging from studies of SARS-CoV-2 [33]. We developed and applied analytical and modelling workflows, using text mining approaches and contents of interaction databases, to propose preliminary insights into COVID-19 molecular mechanisms. The Map continues to grow, following emerging scientific literature. Its content is currently centred on molecular processes involved in SARS-CoV-2 entry and replication, and host-virus interactions. As scientific evidence of host susceptibility, immune response, cell and organ specificity emerge, these will be incorporated into future versions of the Map (Figure 2).

**Figure 2:**
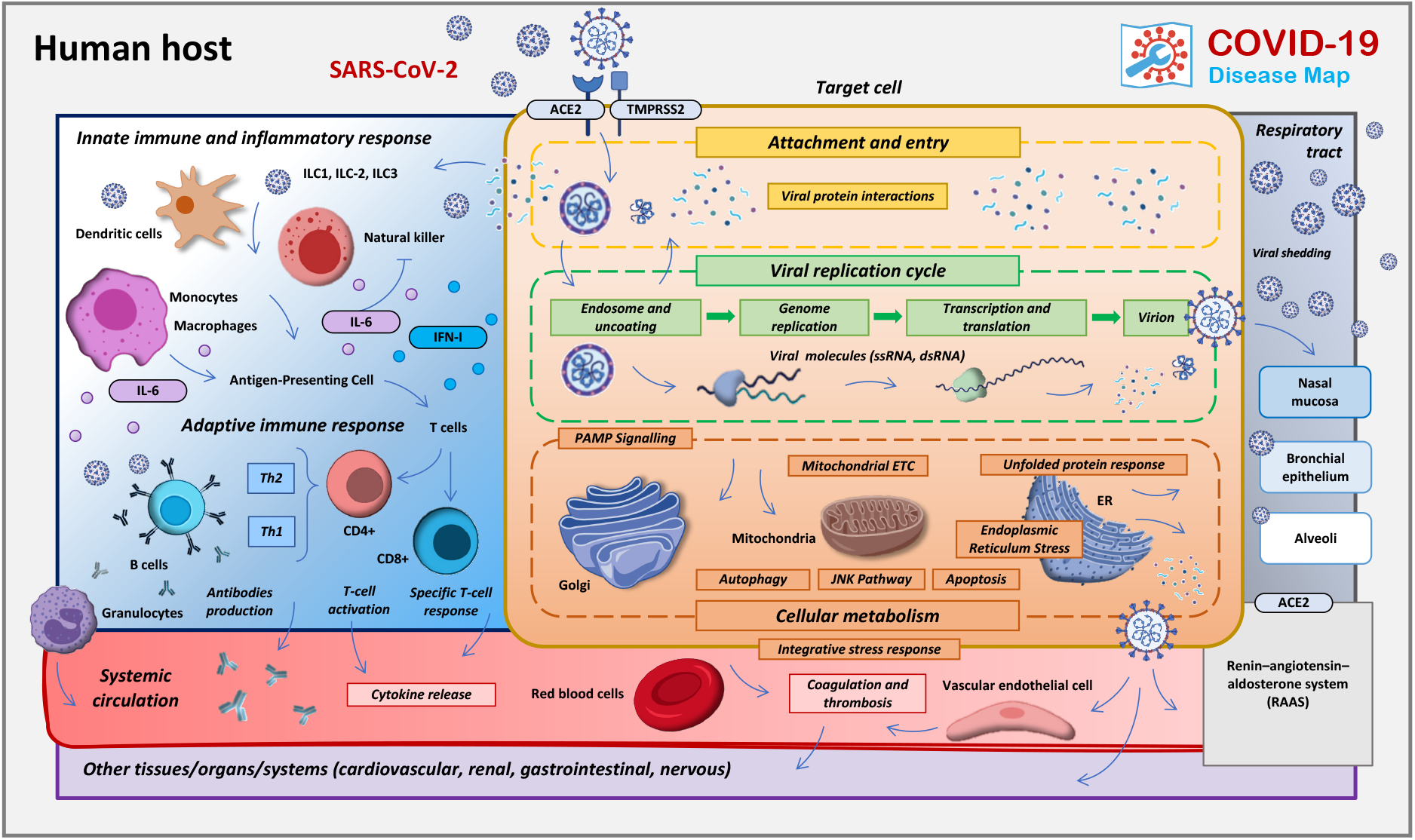
The structure and content of the COVID-19 Disease Map. The areas of focus of the COVID-19 Map biocuration.

While the interactions of SARS-CoV-2 with various host cell types are vital determinants of COVID-19 pathology [2,48–52], the current Map represents an infection of a generic host cell. Several pathways included in the COVID-19 Map are shared between different cell types, for example the IFN-1 pathway found in dendritic and epithelial cells, and in alveolar macrophages [53–57]. Continued annotations of emerging expression data sets and other sources of information will allow the construction of cell-specific versions of the Map to provide an integrated view of the effects of SARS-CoV-2 on the human organism.

SARS-CoV-2 infection and COVID-19 progression are sequential events that start with viral attachment and entry (Figure 3). These events involve various dynamic processes and different time scales that are not captured in static representations of pathways. Correlation of symptoms and potential drugs suggested to date helps downstream data exploration and drug target interpretation in the context of therapeutic interventions.

**Figure 3:**
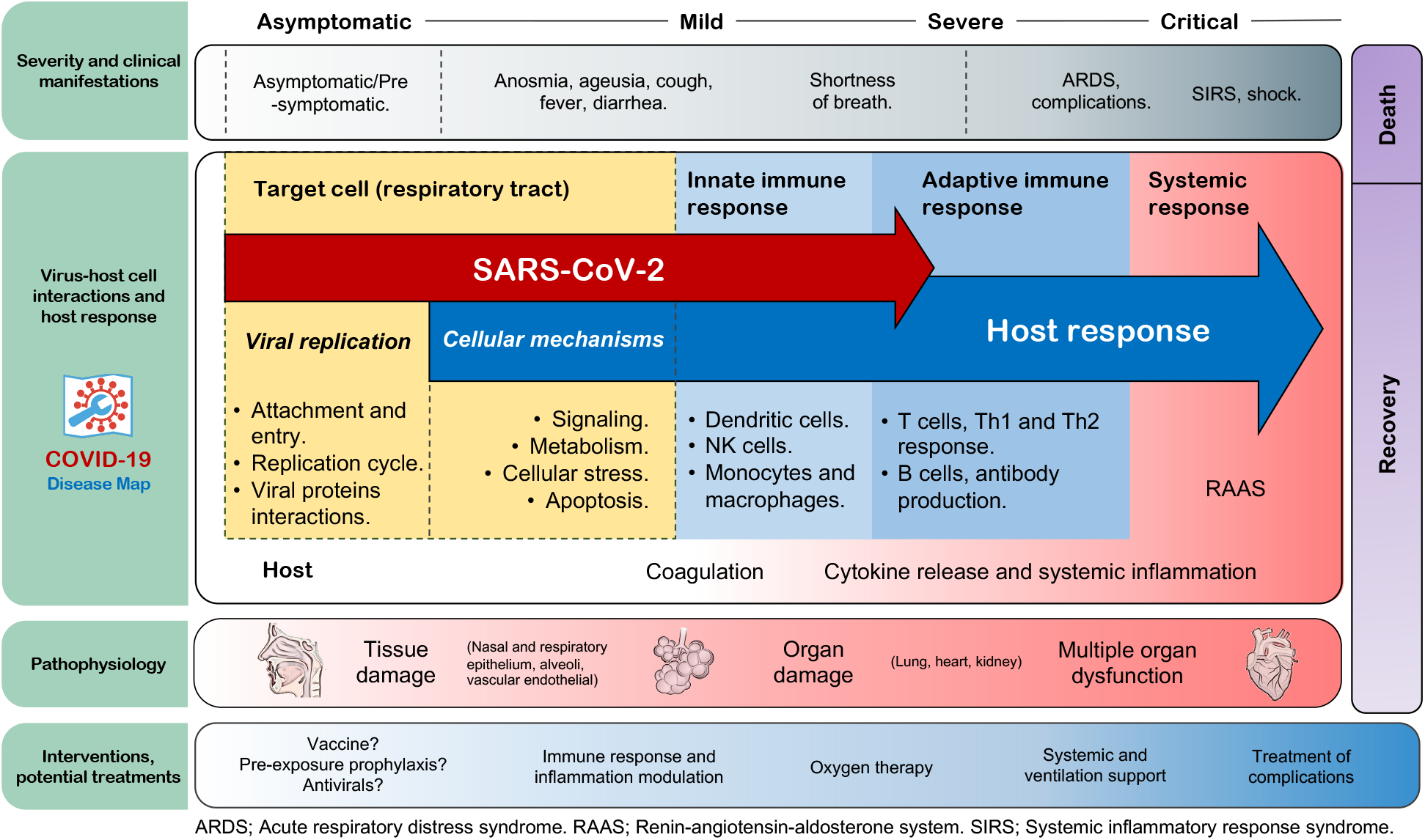
Overview of the Map in the context of COVID-19 progression. Pathways and cell types involved in COVID-19, including some of the most common clinical manifestations and medical management from the moment of infection to disease resolution. The distribution of the elements is for illustrative reference and does not necessarily indicate either a unique/static interplay of these elements or an unvarying progression. For the literature on clinical manifestations see [58–64].

Supplementary Material 1 summarises the contents of the COVID-19 Disease Map diagrams, their central platform of reference. The online version of the table is continuously updated to reflect the evolving content of the COVID-19 Disease Map^24^.

### 3.1 Contents of the Map

#### Virus replication cycle

The virus replication cycle includes binding of the spike surface glycoprotein (S) to angiotensin-converting enzyme 2 (ACE2) mediated by TMPRSS2 [65–68], and other receptors [69,70]. Viral entry occurs either by direct fusion of the virion with the cell membranes or by endocytosis [67,71,72] of the virion membrane, and the subsequent injection of the nucleocapsid into the cytoplasm. Within the host cell, the Map depicts how SARS-CoV-2 hijacks the rough endoplasmic reticulum (RER)-linked host translational machinery for its replication [47,73–78]. The RER-attached translation machinery produces structural proteins, which together with the newly generated viral RNA are assembled into new virions and released to the extracellular space via smooth-walled vesicles [47,73] or hijacked lysosomes [79].

#### Viral subversion of host defence

Endoplasmic reticulum (ER) stress is a consequence of the production of large amounts of viral proteins that create an overload of unfolded proteins [80–82]. The mechanisms of the unfolded protein response (UPR) [83] include the mitigation of the misfolded protein load by increased protein degradation and reduced protein synthesis [84–86]. Malfunctioning proteins and damaged organelles are degraded through the ubiquitin-proteasome system (UPS) and autophagy [87–91]. SARS-CoV-2 may perturb the process of UPS-based protein degradation via the interaction of the Orf10 virus protein with the Cul2 ubiquitin ligase complex and its potential substrates [33,92]. Its involvement in autophagy is less documented [93,94].

This increased burden of misfolded proteins due to viral replication and subversion of mitigation mechanisms may trigger programmed cell death (apoptosis). The Map encodes major signalling pathways triggering this final form of cellular defence against viral replication [95–97]. Many viruses block or delay cell death by expressing anti-apoptotic proteins to maximise the production of viral progeny [98,99], or induce it in selected cell types [97,100–105].

#### Host integrative stress response

SARS-CoV-2 infection damages the epithelium and the pulmonary capillary vascular endothelium [106,107], causing impaired respiratory capacity and leading to acute respiratory distress syndrome (ARDS) in severe forms of COVID-19 [60,108,109]. The release of pro-inflammatory cytokines and hyperinflammation are known complications, causing further widespread damage [110–113]. Coagulation disturbances and thrombosis are associated with severe cases, but unique specific mechanisms have not been described yet [63,114–116]. Nevertheless, it was shown that SARS-CoV-2 disrupts the coagulation cascade and causes renin-angiotensin system (RAS) imbalance [117,118].

ACE2, used by SARS-CoV-2 for host cell entry, is a regulator of RAS and is widely expressed in the affected organs. The diagrams in the repository describe how ACE2-converted angiotensins trigger the counter-regulatory arms of RAS, and the downstream signalling via AGTR1, regulating the coagulation cascade [119–121].

#### Host immune response

The innate immune system detects specific pathogen-associated molecular patterns, through Pattern Recognition Receptors (PRRs), that recognise viral RNA in the endosome during endocytosis, or in the cytoplasm during virus replication. The PRRs activate associated transcription factors promoting the production of antiviral proteins like interferon-alpha, beta and lambda [47,54,55,57,122–127]. SARS-CoV-2 impairs this mechanism [48], but the exact components are yet to be elucidated [128–134]. The Map includes both the virus recognition process and the viral evasion mechanisms. It provides the connection between virus entry, its replication cycle, and the effector pathways of pro-inflammatory cytokines, especially of the interferon type I cascade [2,47,57,130,135–141].

Key metabolic pathways modulate the availability of nutrients and critical metabolites of the immune microenvironment [142]. They are a target of infectious entities that reprogram host metabolism to create favourable conditions for their reproduction [143]. The Map encodes several immunometabolic pathways and provides detailed information about the way SARS-CoV-2 proteins interact with them. The metabolic pathways include heme catabolism [144–146] and its downstream target, the NLRP3 inflammasome [147–152], both affected by SARS-CoV and SARS-CoV-2 proteins [33,153–157], tryptophan-kynurenine metabolism, governing the response to inflammatory cytokines [158–162], and nicotinamide and purine metabolism [163–166] targeted by SARS-CoV-2 [33]. Finally, we represent the pyrimidine synthesis pathway, tightly linked to purine metabolism, affecting viral DNA and RNA synthesis [167–169].

### 3.2 Exploration of the networked knowledge

The pathway diagrams of the COVID-19 Map are constructed by community curators. Their assembly into a repository with standard encoding and annotation, linked to interaction and text mining databases (see Section 2.2) supports exploration to identify crosstalks and functional overlaps across pathways. These analyses allow us to fill gaps in our understanding of COVID-19 mechanisms and generate new testable hypotheses (see Supplementary Material 4). Below, we discuss three examples of our exploration of the networked knowledge in the Map, illustrated in Figure 4.

**Figure 4:**
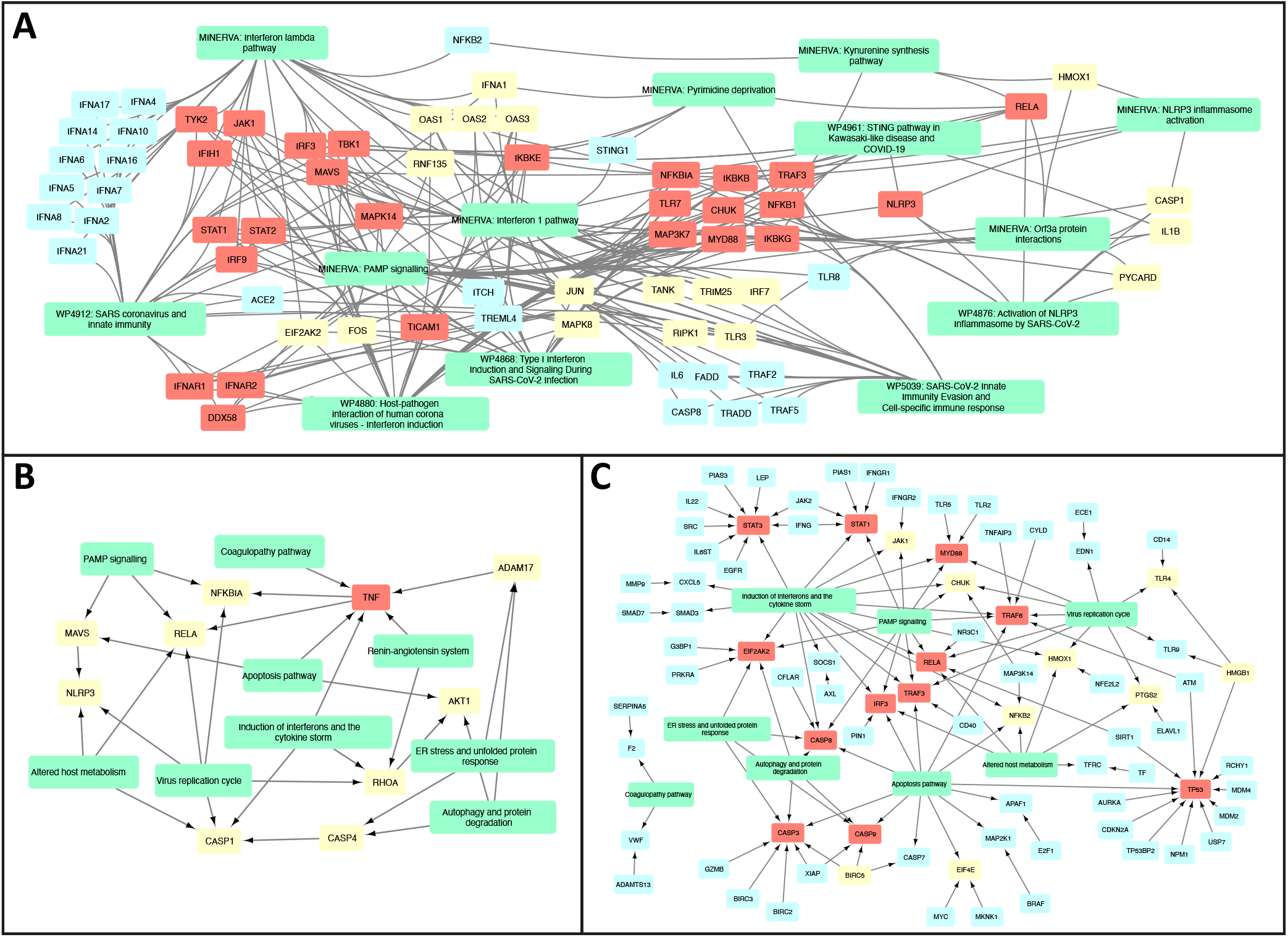
Exploration of the existing and new crosstalks between the diagrams of the COVID-19 Disease Map. The network structure of the diagrams and their interactions based on existing crosstalk (shared elements), new crosstalks and new regulators. A) Existing crosstalks between individual diagrams of IFN-I and RELA-related mechanisms; B) New crosstalks between pathway groups, and C) Novel regulators of existing diagrams as suggested by text mining and interaction databases. Colour code: green - pathways or pathway groups, blue - proteins with two neighbors, yellow - proteins with three or four, red - proteins with five or more. See Supplementary Material 4 for details.

**Figure 4:**
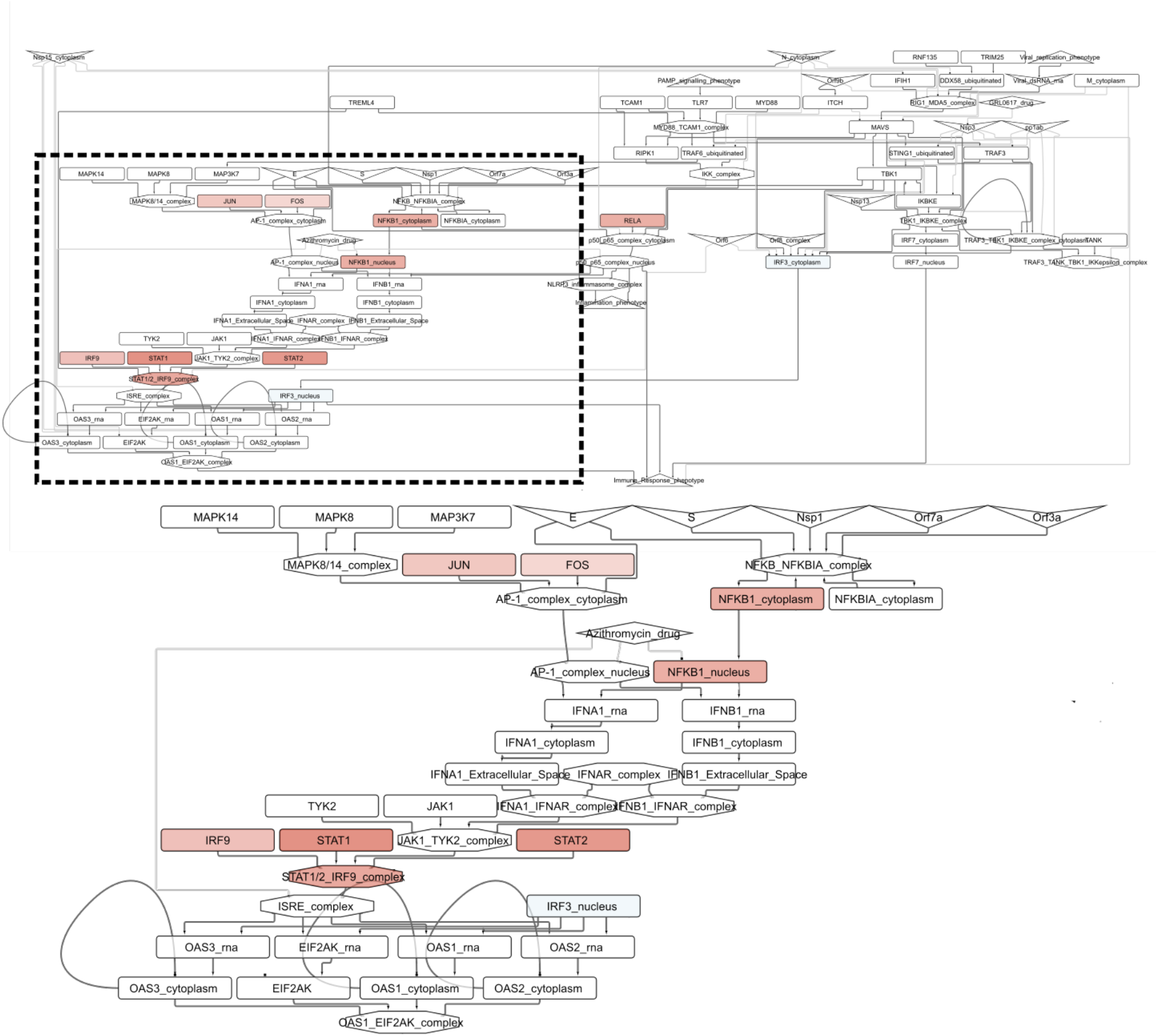
The *Interferon type I signalling pathway* diagram of the COVID-19 Disease Map integrated with TF activity derived from transcriptomics data after SARS-CoV-2 infection. A zoom was applied in the area containing the most active TFs (red nodes) after infection. Node shapes: host genes (rectangles), host molecular complex (octagons), viral proteins (V shape), drugs (diamonds) and phenotypes (triangles).

#### Existing crosstalks between COVID 19 Disease Map diagrams

First, the existing pathway crosstalks emerge when entities are matched between different diagrams (Figure 4A). For instance, they link different pathways involved in type I IFN (IFN-1) signalling. Responses to RNA viruses and pathogen-associated molecular patterns (PAMPs) share common pathways, involving RIG-I/Mda-5, TBK1/IKKE and TLR signalling, leading to the production of IFN-1s, especially IFN-beta [170,171] and IFN-alpha [122]. Downstream, IFN-1 activates Tyk2 and Jak1 protein tyrosine kinases, causing STAT1:STAT2:IRF9 (ISGF3) complex formation to promote the transcription of IFN-stimulated genes (ISGs). Importantly, TBK1 also phosphorylates IKBA, an NF-kB inhibitor, for proteasomal degradation in crosstalk with the UPS pathway, allowing free NF-kB and IRF3 to co-activate ISGs [172]. Another TBK1 activator, STING, links IFN signalling with pyrimidine metabolism.

SARS-CoV-2 M protein affects these IFN responses by inhibiting the RIG-I:MAVS:TRAF3 complex and TBK1, preventing IRF3 phosphorylation, nuclear translocation, and activation [173]. In severe COVID-19 cases, elevated NF-kB activation, associated with impaired IFN-1 [54] may be a host attempt to compensate for the lack of IFN-1 activation [174], leading to NF-kB hyperactivation and release of pro-inflammatory cytokines. Moreover, SARS-CoV-1 viral papain-like-proteases, contained within the nsp3 and nsp16 proteins, inhibit STING and its downstream IFN secretion [175]. Defective responses involving these pathways and other regulatory factors may impair the IFN response against SARS-CoV-2, and explain persistent blood viral load and an exacerbated inflammatory response in COVID-19 patients [54].

#### New crosstalks from interaction and text mining datasets

New relationships emerging from associated interaction and text mining databases (see Section 2.2) suggest new pathway crosstalks (see Figure 4B). One of these is the interaction of ER stress and the immune pathways, as PPP1R15A regulates the expression of TNF and the translational inhibition of both IFN-1 and IL-6 [176]. This finding coincides with the proposed interaction of pathways responsible for protein degradation and viral detection, as SQSTM1, an autophagy receptor and NFKB1 regulator, controls the activity of cGAS, a double-stranded DNA detector [177,178]. Another association discovered in text mining data is ADAM17 and TNF release from the immune cells in response to ACE2-S protein interaction with SARS-CoV-1 [179], potentially increasing the risk of COVID-19 infection [180]. This new interaction connects diagrams of the i) “Viral replication cycle” via ACE2-S protein interactions, ii) “Viral subversion of host defence mechanisms” via ER stress, iii) “Host integrative stress response” via the renin-angiotensin system and iv) “Host innate immune response” via pathways implicating TNF signalling.

#### Novel regulators of key pathway proteins

Finally, using interaction and text mining databases, we can identify potential novel regulators of proteins in the Map (see Figure 4C). These proteins take no part in the current version of the Map but interact with molecules already represented in at least one of the diagrams. An example of such a novel regulator is NFE2L2, which controls the activity of HMOX1 in the context of viral infection [181,182]. In turn, HMOX1 controls immunomodulatory heme metabolism [144,145] and mechanisms of viral replication [183] and is a target of SARS-CoV-2 Orf3a protein [157,183]. The suggested NFE2L2-HMOX1 interaction is supported by the literature reports of NFE2L2 importance in COVID-19 cardiovascular complications due to crosstalk with the renin-angiotensin signalling pathway [184] and potential interactions with viral entry mechanisms [185]. Interestingly, the modulation of the NFE2L2-HMOX1 axis was already proposed as a therapeutic measure [186], making it an interesting extension of the COVID-19 Disease Map.

### 3.3 Biocuration roadmap

COVID-19 Disease Map pathways span a range of currently known host-cell virus interactions and mechanisms. Nevertheless, certain aspects of the disease are not represented in detail, particularly cell-type-specific immune response, and susceptibility features. Their mechanistic description is of great importance, as suggested by clinical reports on the involvement of these pathways in the molecular pathophysiology of the disease. The mechanisms outlined below will be the next targets in our curation roadmap.

#### Cell type-specific immune response

COVID-19 causes serious disbalance in multiple populations of immune cells, including peripheral CD4+ and CD8+ cytotoxic T lymphocytes, B cells and NK cells [111,162,187–190]. This may be the result of functional exhaustion due to SARS-CoV-2 S protein and excessive pro-inflammatory cytokine response [188,191], promoted by an abnormal increase of the Th17:Treg cell ratio [192]. Moreover, the ratio of naive-to-memory helper T-cells increases while the level of T regulatory cells decreases in severe cases [193]. Pulmonary recruitment of lymphocytes into the airways, including Th17 and cytotoxic CD8+ T-cells [194], may explain this imbalance and the increased neutrophil-lymphocyte ratio in peripheral blood [187,195,196]. To address this aspect of the disease we plan to implement cell type representations of different populations, and encode their cell surface receptors and transition mechanisms. With the help of single-cell omics profiling, we plan to adapt these to reflect COVID-19 specificity.

#### Susceptibility features of the host

SARS-CoV-2 infection is associated with increased morbidity and mortality in individuals with underlying medical conditions, chronic diseases or a compromised immune system [197–200]. Groups at risk are men, pregnant and postpartum women, and individuals with high occupational viral exposure [201–203]. Other susceptibility factors include the ABO blood groups [204–211] and respiratory conditions [212–217].

Importantly, age is one of the key aspects contributing to the severity of the disease [199,218]. Age-related elevated levels of inflammation [218–221], immunosenescence and cellular stress of ageing cells [108,199,218,222,223] may contribute to the risk. In contrast, children are generally less likely to develop severe disease [224,225], with the exception of infants [108,226–228]. However, some previously healthy children and adolescents can develop a multisystem inflammatory syndrome following SARS-CoV-2 infection [229–233].

Finally, several genetic factors have been proposed and identified to influence susceptibility and severity, including the ACE2 gene, HLA locus, errors influencing type I IFN production, TLR pathways, myeloid compartments, as well as cytokine polymorphisms [207,234–241].

Connecting the susceptibility features to specific molecular mechanisms will allow us to better understand the contributing factors. These features can be directly incorporated as elements of relevant diagrams. Another possibility is connecting the diagrams of the Map to clinical and phenotypic data following big data workflows as demonstrated in other settings [3,242]. This can lead to a series of testable hypotheses, including the role of lipidomic reprogramming [243,244] or of vitamin D [245–247] in modifying the severity of the disease. Another testable hypothesis is that the immune phenotype associated with asthma inhibits pro-inflammatory cytokine production and modifies gene expression in the airway epithelium, protecting against severe COVID-19 [216,217,248].

## 4. Bioinformatics analysis and computational modelling roadmap for hypothesis generation

To understand complex and often indirect dependencies between different pathways and molecules, we need to combine computational and data-driven analyses. Standardised representation and programmatic access to the contents of the COVID-19 Disease Map support reproducible analytical and modelling workflows. Here, we discuss the range of possible approaches and demonstrate preliminary results, focusing on interoperability, reproducibility, and applicability of the methods and tools.

### 4.1 Data integration and network analysis

Visualisation of omics datasets can help contextualise the Map with experimental data, creating data-specific blueprints. They can highlight parts of the Map that are active in one condition versus another. Combining information contained in multiple omics platforms can make patient stratification more powerful, by reducing the number of samples needed or by augmenting the precision of the patient groups [249,250]. Approaches that integrate multiple data types without the accompanying mechanistic diagrams [251–253] produce patient groupings that are difficult to interpret. In turn, classical pathway analyses often produce long lists mixing generic and cell-specific pathways, making it challenging to pinpoint relevant information. Using disease maps to interpret omics-based clusters addresses the issues related to contextualised visual data analytics.

#### Footprint based analysis

Footprints are signatures of a molecular regulator determined by the expression levels of its targets [254]. Combining multiple omics readouts and multiple measurements can increase the robustness of such signatures. Nevertheless, an essential component is the mechanistic description of the targets of a given regulator, allowing computation of its footprint. With available SARS-CoV-2 related omics and interaction datasets [255], it is possible to infer which TFs and signalling pathways are affected upon infection [256]. Combining the COVID-19 Disease map regulatory interactions with curated collections of TF-target interactions like DoRothEA [257] will provide a contextualised evaluation of the effect of SARS-CoV-2 infection at the TF level.

#### Virus–host interactome

The virus–host interactome is a network of viral-human protein-protein interactions (PPIs) that can help to understand the mechanisms of viral diseases [33,258–260]. It can be expanded by merging virus-host PPI data with human PPI and protein data [261] to discover clusters of interactions indicating human mechanisms and pathways affected by the virus [262]. These clusters can be interpreted at the mechanistic level by visual exploration of COVID-19 Disease Map diagrams. In addition, these clusters can potentially reveal additional pathways to add to the COVID-19 Disease Map (e.g. E protein interactions or TGF beta diagrams) or suggest new interactions to introduce into the existing diagrams.

### 4.2 Mechanistic and dynamic computational modelling

Computational modelling is a powerful approach that enables *in silico* experiments, produces testable hypotheses, helps elucidate regulation and, finally, can suggest via predictions novel therapeutic targets and candidates for drug repurposing.

#### Mechanistic pathway modelling

Molecular interactions of a given pathway can be coupled with its endpoint and contextualised using omics datasets. For instance, HiPathia uses transcriptomic or genomic data to estimate the functional profiles of a pathway in relation to their endpoints of interest [263,264]. Such mechanistic modelling can be used to predict the effect of interventions,, for example effects of drugs on their targets [265]. HiPathia integrates directly with the diagrams of the COVID-19 Map using the SIF format provided by CaSQ (see Section 2.3), as well as with the associated interaction databases (see Section 2.2). The drawback of such approaches is their computational complexity, limiting the size of the diagrams they can process. Large-scale mechanistic pathway modelling requires their transformation into causal networks. CARNIVAL [266] combines the causal representation of networks [12] with transcriptomics, (phospho)proteomics, or metabolomics data [254] to contextualise cellular networks and extract mechanistic hypotheses. The algorithm identifies a set of coherent causal links connecting upstream drivers such as stimulations or mutations to downstream changes in transcription factor activities.

#### Discrete computational modelling

Discrete modelling allows analysis of the dynamics of molecular networks to understand their complexity under disease-related perturbations. COVID-19 Disease Map diagrams, translated to SBML qual using CaSQ (see Section 2.3), can be directly imported by tools like Cell Collective [267] or GINsim [45] for analysis. Cell Collective is an online modelling platform^25^ that provides features for real-time simulations and analysis of complex signalling networks. References and layout are used for model visualisation, supporting the interpretation of the results. In turn, GINsim provides a range of analysis methods, including identification of the states of convergence of a given model (attractors). Model reduction functionality can also be employed to facilitate the analysis of large-scale models.

#### Multiscale and stochastic computational modelling

Viral infection and immune response are processes that span many scales, from molecular interactions to multicellular behaviour. Modelling of such complex scenarios requires a multiscale computational architecture, where single cell models run in parallel to capture behaviour of heterogeneous cell populations and their intercellular communications. Multiscale agent-based models offer such architecture, and can simulate processes at different time scales, e.g. diffusion, cell mechanics, cell cycle, or signal transduction [268,269]. An example of such approach is PhysiBoSS [270], which combines the computational framework of PhysiCell [271] with MaBoSS [272], a tool for stochastic simulations of logical models to study of transient effects and perturbations [273]. Implementing detailed COVID-19 signalling models in the PhysiBoSS framework may help to better understand complex dynamics of interactions between immune system components and the host cell.

### 4.3 Case study: RNA-Seq-based analysis of transcription factor activity

We measured the effect of COVID-19 at the transcription factor (TF) activity level by applying VIPER [274] combined with DoRothEA regulons [257] on RNA-seq datasets of the SARS-CoV-2 infected Calu-3 cell line [126]. Then, we mapped the TFs normalised enrichment score (NES) on the *Interferon type I signalling pathway* diagram of the COVID-19 Disease Map using the SIF files generated by CaSQ (see Section 2.3). As highlighted in Figure 4, our manually curated pathway included some of the most active TFs after SARS-CoV-2 infection, such as STAT1, STAT2, IRF9 and NFKB1. These are well known components of cytokine signalling and antiviral responses [275,276]. Interestingly, they are located downstream of various viral proteins (*E, S, Nsp1, Orf7a* and *Orf3a*) and members of the MAPK pathway (*MAPK8, MAPK14* and *MAP3K7*). SARS-CoV-2 infection is known to promote MAPK activation, which mediates the cellular response to pathogenic infection and promotes the production of pro-inflammatory cytokines [255]. These conclusions can be used to investigate response of the human cells to SARS-CoV-2 infection.

### 4.4 Case study: RNA-seq-based analysis of pathway signalling

The Hipathia [263] algorithm was used to calculate the level of activity of the subpathways from the COVID-19 Apoptosis diagram. We used a public RNA-seq dataset from human SARS-CoV-2 infected lung cells (GEO GSE147507). We treated the RNA-seq gene expression data with the Trimmed Mean of M values (TMM) normalisation [277], rescaled to range [0;1] for the calculation of the signal and normalised using quantile normalisation [278]. Using the normalised gene expression values we calculated the level of activation of the subpathways, then we used case/control contrast with a Wilcoxon test to assess differences in signalling activity between the two conditions.

Results of the Apoptosis pathway analysis can be seen in Figure 5 and Supplementary Material 5. The analysis shows an overactivation of several circuits (series of causally connected elements), specifically upstream of the effector protein BAX, led by the overexpression of the BAD protein, inhibiting BCL2-MCL1-BCL2L1 complex, which in turn inhibits BAX. Indeed, SARS-CoV-2 infection can invoke caspase8-induced apoptosis [279], where BAX together with the ripoptosome/caspase-8 complex, may act as a pro-inflammatory checkpoint [280]. This result is supported by studies in SARS-CoV-1, showing BAX overexpression following infection [104,281]. Overall, our findings recapitulate reported outcomes and suggest that with evolving contents of the COVID-19 Disease Map and new transcriptomic data becoming available, new mechanism-based hypotheses can be formulated.

**Figure 5.**
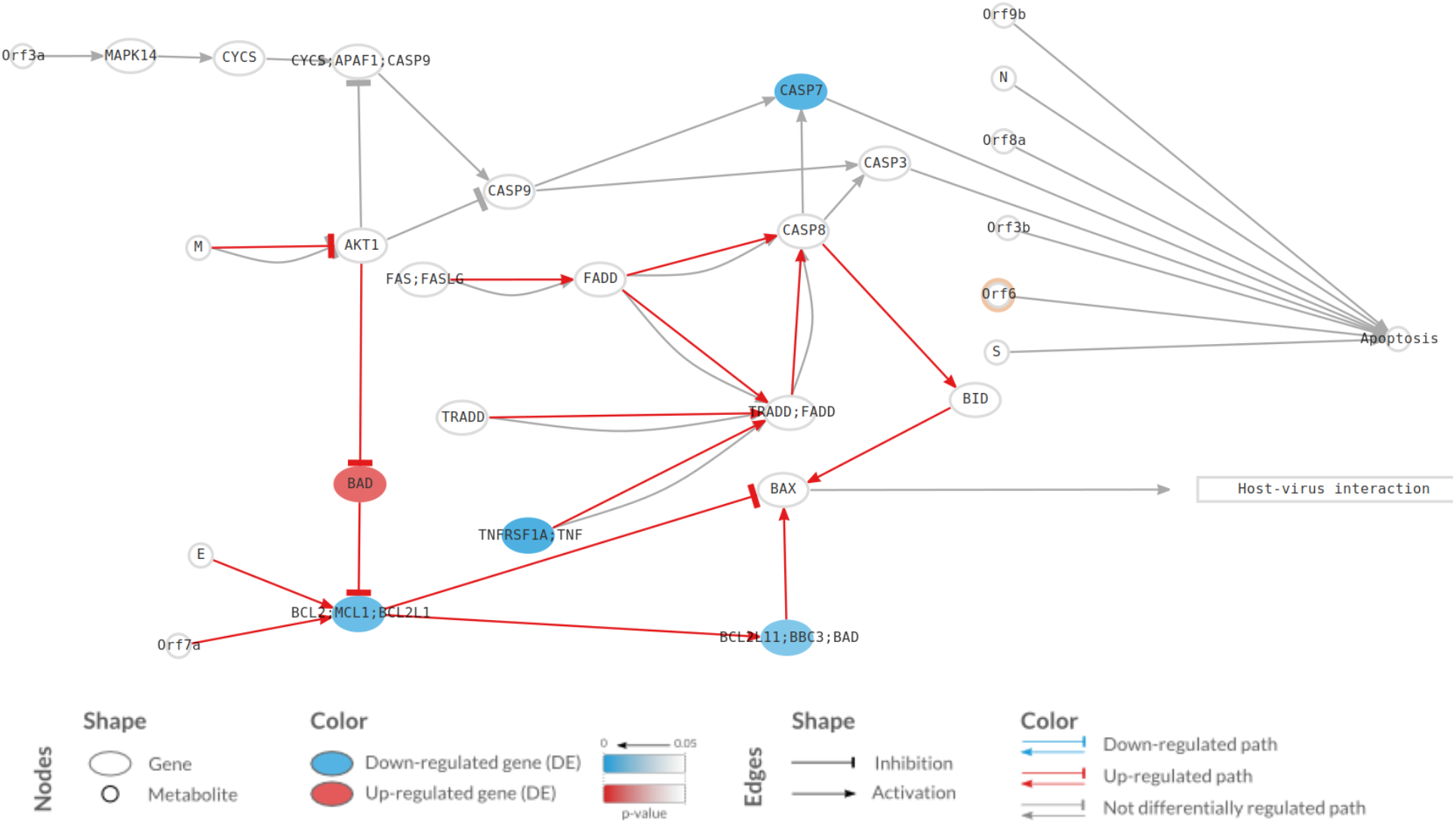
Representation of the activation level of Apoptosis pathway in SARS-CoV-2 infected lung cell lines. Activation levels were calculated using transcriptional data from GSE147507 and the Hipathia mechanistic pathway analysis algorithm. Each node represents a gene (ellipse), a metabolite (circle) or a function (square). The pathway is composed of circuits from a receptor gene/metabolite to an effector gene/function, which take into account interactions simplified to inhibitions or activations (see Section 2.3, SIF format). Significantly deregulated circuits are highlighted by color arrows (red: activated in infected cells). The color of the node corresponds to the level of differential expression in SARS-CoV-2 infected cells vs normal lung cells. Blue: down-regulated elements, red: up-regulated elements, white: elements with no statistically significant differential expression.

### 4.5 Parallel efforts

There are parallel efforts towards modelling of COVID-19 mechanisms, providing a complementary source of information and their future integration will create an even broader toolset to tackle the pandemic.

The modified Edinburgh Pathway Notation (mEPN) [282] is a scheme for visual encoding of molecular processes in diagrams that also function as Petri nets, allowing activity simulations using the BioLayout tool [283]. The current mEPN COVID-19 model details the replication cycle of SARS-CoV-2, integrated with a range of host defence systems. Currently, models constructed in mEPN can be translated to SBGNML, but without the information related to their function as Petri nets.

The COVID-19 Disease Map can also support kinetic modelling to quantify the behaviour of pathways and evaluate the dynamic effects of perturbations. However, it is necessary to assign a kinetic equation or a rate law to every reaction in the diagram to be analysed. This process is challenging and requires support of tools like SBMLsqueezer [20] and reaction kinetics databases like SABIO-RK [284]. Nevertheless, the most critical factor is the availability of experimentally validated parameters that can be reliably applied in SARS-CoV-2 modelling scenarios.

## 5. Discussion

COVID-19 literature is growing at great speed, fueled by global research efforts to investigate the pathophysiology of SARS-CoV-2 infection and to better understand susceptibility factors and identify molecular targets of therapeutic intervention. We need to improve the use of this knowledge by tools and approaches to extract, formalise and integrate relevant information, and by application of analytical frameworks to generate testable hypotheses from systems level models.

The COVID-19 Disease Map is an open access knowledgebase and computational repository. On the one hand, it is a graphical, interactive representation of disease-relevant molecular mechanisms linking many knowledge sources. On the other hand, it is a computational resource of curated content for graph-based analyses and disease modelling. It offers a shared mental map for understanding the dynamic nature of the disease at the molecular level and also its dynamic propagation at a systemic level. Thus, it provides a platform for a precise formulation of models, accurate data interpretation, monitoring of therapy, and potential for drug repositioning.

The COVID-19 Disease Map diagrams describe molecular mechanisms of COVID-19. These diagrams are grounded in the relevant published SARS-CoV-2 research, completed where necessary by mechanisms discovered in related beta-coronaviruses. With an unprecedented effort of community-driven biocuration, over forty diagrams with molecular resolution were constructed since March 2020, shared across three platforms.

This large community effort shows that expertise in biocuration, clear guidelines and text mining solutions can accelerate the passage from data generated in the published literature to a meaningful mechanistic representation of knowledge. This exercise in quick research data generation and knowledge accumulation may serve as a blueprint for a formalised and standardised streamline of well-defined tasks.

Moreover, by developing reproducible analysis pipelines for the contents of the Map we promote early harmonisation of formats, support of standards, and transparency in all steps. Preliminary results of such efforts are illustrated in case studies above. Importantly, biocurators and domain experts participate in the analysis, helping to evaluate the outcomes and correct the curated content if necessary. This way, we improve the quality of the analysis and increase reliability of the models in generating useful predictions.

This approach to an emerging pandemic leveraged the capacity and expertise of an entire swath of the bioinformatics community, bringing them together to improve the way we build and share knowledge. By aligning our efforts, we strive to provide COVID-19 specific pathway models, synchronise content with similar resources and encourage discussion and feedback at every stage of the curation process.

The COVID-19 Disease Map community is open and expanding as more people with complementary expertise join forces. In the longer run, the COVID 19 Disease Map content will be used to facilitate the finding of robust signatures related to SARS-CoV-2 infection predisposition or response to various treatments, along with the prioritization of new potential drug targets or drug candidates. The project aims to provide the tools to deepen our understanding of the mechanisms driving the infection and help boost drug development supported by testable suggestions. Such an approach may help dealing with new waves of COVID-19 or similar pandemics in the long-term perspective.

## Authors’ contributions

M. Ostaszewski, A. Niarakis, A. Mazein and I. Kuperstein planned and coordinated the project;

R. Phair, A. Orta-Resendiz, J-M. Ravel, R. Fraser, V. Ortseifen and S. Marchesi advised the project as domain experts;

M. Ostaszewski, A. Niarakis, A. Mazein, I. Kuperstein, V. Singh, S.S. Aghamiri, M.L. Acencio, E. Glaab, A. Ruepp, G. Fobo, C. Montrone, B. Brauner, G. Frischman, L.C.M. Gomez, J. Sommers, M. Hoch, S. Gupta, J. Scheel, H. Borlinghaus, T. Czauderna, F. Schreiber, A. Montagud, M. Ponce de Leon, A. Funahashi, Y. Hiki, N. Hiroi, T.G, Yamada, A. Dräger, A. Renz, M. Naveez, Z. Bocskei, F. Messina, D. Börnigen, L. Fergusson, M. Conti, M. Rameil, V. Nakonecnij, J. Vanhoefer, L. Schmiester, M. Wang, E.E. Ackerman, J. Shoemaker, J. Zucker, K. Oxford, J. Teuton, E. Kocakaya, G.Y. Summak, K. Hanspers, M. Kutmon, S. Coort, L. Eijssen, F. Erhart, D.A.B. Rex, D. Slenter, M. Martens, N. Pham, R. Haw, B. Jassal, L. Matthews, M. Orlic-Milacic, A.S. Ribeiro, K. Rothfels, V. Shamovsky, R. Stephan, C. Sevilla and T. Varusai curated and reviewed the diagrams;

M. Ostaszewski, P. Gawron, E. Smula, L. Heirendt, V. Satagopam, G. Wu, A. Riutta, M. Golebiewski, S. Owen, C. Goble and X. Hu designed, developed and implemented key elements of the data sharing and communication infrastructure;

RW. Overall, D. Maier, A. Bauch, B.M. Gyori, J.A. Bauch, C. Vega, V. Groues, M. Vazquez, P. Porras, L. Licata, M. Ianucelli, F. Sacco, A. Nestorova, A. Yuryev, A. de Waard designed and developed the contents of interaction and pathway databases, and text mining platforms and their visualisation and interoperability functionalities;

A. Niarakis, D. Turei, A. Luna, O. Babur, S. Soliman, A. Valdeolivas, M. Esteban, M. Pena, K. Rian, T. Helikar, B.L. Puniya, D. Modos, A. Treveil, M. Olbei, B. De Meulder, A. Dugourd, A. Naldi, V. Noel, L. Calzone developed format interoperability, analysis and modelling workflows;

C. Sander, E. Demir, T. Korcsmaros, T. Freeman, F. Augé, J.S. Beckmann, J. Hasenauer, O. Wolkenhauer, E.L. Wilighagen, A.R. Pico, C.T. Evelo, M.E. Gillespie, L. Stein, H. Hermjakob, P. D’Eustachio, J. Saez-Rodriguez, J. Dopazo, A. Valencia, H. Kitano, E. Barillot, C. Auffray, R. Balling, R. Schneider defined the strategy and scope of the project and revised its progress;

M. Ostaszewski, A. Niarakis, A. Mazein and I. Kuperstein wrote the manuscript;

A. Orta-Resendiz, I. Kuperstein and A. Mazein designed the overview figures;

A. de Waard, P. D’Eustachio, J.S. Beckmann and L.D. Stein revised and contributed significantly to the structure of the manuscript;

All authors have revised, read and accepted the manuscript in its final form.

## Acknowledgements

We would like to thank Andjela Tatarovic, architect, and Gina Crovetto, researcher in the field of cancer, for their help with the design of the top-level view diagrams. We would like to acknowledge the Responsible and Reproducible Research (R3) team of the Luxembourg Centre for Systems Biomedicine for supporting the project and providing necessary communication and data sharing resources. The work presented in this paper was carried out using the ELIXIR Luxembourg tools and services. EG acknowledges support from the Luxembourg National Research Fund (FNR) as part of the COVID-19 Fast-Track grant programme (COVID-19/2020-1/14715687/CovScreen). AM, MPL, MV and AV would like to acknowledge two European Commission grants: INFORE (H2020-ICT-825070) and PerMedCoE (H2020-ICT-951773).

## Supplementary Materials

Supplementary Material 1: COVID-19 Disease Map diagrams

Supplementary Material 2: Biocuration platforms and formats

Supplementary Material 3: Description of bioinformatic resources

Supplementary Material 4: Exploration of crosstalks in the COVID-19 Disease Map diagrams

Supplementary Material 5: Results of the Hipathia analysis of the Apoptosis pathway

## Footnotes

1 https://covid19.who.int/

2 https://covid.pages.uni.lu/map_contents

3 https://memote.io

4 https://fairdomhub.org/documents/661

5 https://reactome.org/community/training

6 https://www.wikipathways.org/index.php/Help:Editing_Pathways

7 http://celldesigner.org

8 https://newteditor.org

9 https://github.com/sbgn/ySBGN

10 https://www.semanticscholar.org/cord19/download (accessed on 20.10.2020)

11 https://data.mendeley.com/datasets/h9vs5s8fz2/draft?a=f40961bb-9798-4fd1-8025-e2a3ba47b02e

12 https://www.ebi.ac.uk/intact/imex/main.xhtml?query=annot:“dataset:coronavirus”

13 https://signor.uniroma2.it/covid/

14 https://pathwaystudio.com

15 https://biokb.lcsb.uni.lu

16 https://ailani.ai

17 https://git-r3lab.uni.lu/covid/models/-/tree/master/Resources/Text%20mining

18 https://opennlp.apache.org

19 https://rupertoverall.net/covidminer

20 https://github.com/golovatenkop/rnef2sbgn

21 here, SBML stands for two formats: CellDesigner SBML and SBML with *layout* and *render* packages

22 http://www.ebi.ac.uk/sbo/main/

23 http://www.cbmc.it/fastcent/doc/SifFormat.htm

24 https://covid.pages.uni.lu/map_contents

25 https://cellcollective.org

