## Supplementary Table 1 for "COVID-19 Disease Map, a computational knowledge repository of SARS-CoV-2 virus-host interaction mechanisms"

### Supplementary Table 1: COVID-19 Disease Map diagrams

The Table summarises the main diagrams of the COVID-19 Disease Map Project, their scope and direct link. HCoV - Human Coronavirus. Online version at https://covid.pages.uni.lu/map_contents

| Pathway | Diagrams: Resource | Diagrams: Access | Virus proteins |
|---|---|---|---|
| **Virus replication cycle:** Attachment and entry | WikiPathways | WP4846 | All |
| | Reactome | R-HSA-9678110, R-HSA-9694614 | |
| | MINERVA (gitlab) | Virus replication cycle (gitlab) | |
| **Virus replication cycle:** Transcription, translation and replication | WikiPathways | WP4846 | All |
| | Reactome | R-HSA-9679504, R-HSA-9694676<br>R-HSA-9679514, R-HSA-9694682<br>R-HSA-9683701, R-HSA-9694635 | |
| | MINERVA (gitlab) | Virus replication cycle (gitlab)<br>RTC and transcription (gitlab)<br>Nsp9 interactions (gitlab)<br>E protein interactions (gitlab) | |
| **Virus replication cycle:** Assembly and release | Reactome | R-HSA-9679509, R-HSA-9694322 | All |
| | MINERVA (gitlab) | Virus replication cycle (gitlab)<br>Nsp4 and Nsp6 interactions (gitlab) | |
| **Viral subversion of host defence:** ER stress and unfolded protein response | WikiPathways | WP4861 | S (HCoV OC43)<br>4a (MERS)<br>Orf8ab, E, nsp15 (SARS-CoV-1) |
| | MINERVA (gitlab) | ER stress (gitlab) | |
| **Viral subversion of host defence:** Autophagy and protein degradation | WikiPathways | WP4860, WP4936, WP4863 | nsp6 (SARS-CoV-1)<br>nsp567 (PRRSV)<br>Nsp6, Orf3, Orf10 (SARS-CoV-2) |
| | MINERVA (gitlab) | Orf10 Cul2 pathway (gitlab) | |
| **Viral subversion of host defence:** Apoptosis | WikiPathways | WP4864 | Orf3a, Orf3b, Orf6, Orf8a, Orf7a, Orf9b, E, M, N, S (SARS-CoV-1)<br>Nsp4. Nsp5, Nsp6. Nsp7, Nsp8, Orf9c (SARS-CoV-2) |
| | MINERVA (gitlab) | Apoptosis pathway (gitlab)<br>JNK pathway (gitlab)<br>ETC disruption (gitlab) | |
| **Integrative stress response:** Renin-angiotensin system | WikiPathways: | WP4883, WP4799, WP4965 | S (SARS-CoV-2) |
| | MINERVA (gitlab) | Renin-angiotensin pathway (gitlab) | |
| **Integrative stress response:** Coagulopathy | WikiPathways | WP4927 | S (SARS-CoV-2) |
| | MINERVA (gitlab) | Coagulation pathway (gitlab) | |
| **Innate Immune Response:** PAMP signalling | WikiPathways | WP4912 | nsp3, nsp15, Orf3b, Orf8, Orf9, M, N, S (SARS-CoV-1) |
| | MINERVA | PAMP signalling (gitlab) | |
| **Innate Immune Response:** Induction of interferons and the cytokine storm | WikiPathways | WP4868, WP4880, WP4876 | 4a, 4b, PLPro, S (MERS)<br>nsp3, nsp13, Orf3a, Orf6, Orf7a, Orfab, Orfb, M, N, S (SARS-CoV-1) |
| | MINERVA | Interferon I pathway | |
| **Innate Immune Response:** Altered host metabolism | WikiPathways | WP4853 | Nsp14, Orf3a, E, M, N, S (SARS-CoV-2) |
| | MINERVA (gitlab) | Heme Oxygenase pathway (gitlab)<br>Kynurenine synthesis pathway (gitlab)<br>Amino sugar and nucleotide sugar metabolism (gitlab)<br>Pyrimidine deprivation pathway (gitlab) | |
