## Supplementary Material 2 for "COVID-19 Disease Map, a computational knowledge repository of SARS-CoV-2 virus-host interaction mechanisms"

### Supplementary Material 2:
### Biocuration platforms and formats

#### Biocuration platforms

##### Individual diagrams

Individual diagrams, encoded in systems biology layout-aware formats (see below) are developed by several separate biocuration groups following an established approach [PMID:26235082; PMID:23832570; PMID:26192618; PMID:32316560; PMID:32311035; PMID:30520978; PMID:29726961], previously used for influenza viral infection [PMID:24088197], respiratory disorders [PMID:30133857], inflammation [PMID:32893032] or rheumatoid arthritis [PMID:32311035]. Such a community-based curation is a challenge and requires intensive communication. Thus, we shared curation topics, e.g., relevant pathways or particular SARS-CoV-2 proteins across the community to cover the available literature better and identify synergies. We established curation guidelines[1] to ensure proper representation and annotation of the crucial features of the diagrams. Notably, we perform regular technical reviews of the diagrams following a previously established checklist to harmonise their content.

Diagrams developed by independent curation groups have a single visualisation endpoint provided by the MINERVA Platform [PMID:28725475]. The platform integrates data from drug, chemical, and miRNA target databases, enabling systematic identification of entities targeted by these molecules. Diagrams are accessible as submaps and are cross-searchable, such that queries for elements or targets return results for all submaps simultaneously. The entry-level view is based on the synthesis of illustrations discussed in Section 3 (see Figure 2), giving users intuitive access to respective submaps.

[1] https://docs.google.com/document/d/1DFfJZe2xjXrKMHorp_-7hlqWoNVSmx6sIzAaRzXgtQs

#### Reactome

Reactome is an open-source, open-access, manually curated, and peer-reviewed pathway database [PMID:31691815], organising molecules and their functional relations into biological pathways. Biocuration efforts in Reactome initially focused on SARS-CoV-1 as its proteins and their functions are extensively documented in the experimental literature. Reactome curators were assigned a sub-pathway from the viral life cycle, a host pathway, or potential therapeutics. Curators were supported by an editorial manager and a dedicated SARS literature triage process. This effort created a set of pathways for SARS-CoV-1, which in turn provided the basis for computational inference [PMID:17367534] of the corresponding SARS-CoV-2 pathways based on structural and functional homologies between the two viruses. The computationally inferred SARS-CoV-2 infection pathway events and entities were then reviewed and manually curated using published SARS-CoV-2 experimental data.

#### WikiPathways

WikiPathways is a community-curated, open-source, and open-access pathway database. Building on the MediaWiki platform, the central idea of WikiPathways is the engagement of the scientific community in the curation of biological pathway models. The open-source pathway editor PathVisio [PMID:25706687] enables researchers to curate pathway models with detailed annotation of pathway elements through the integrated BridgeDb identifier mapping framework [PMID:20047655]. All pathways are stored in the Graphical Pathway Markup Language (GPML), integrating detailed annotations of the data nodes and interactions with graphical layout information. WikiPathways COVID-19 content is available via a dedicated pathway portal[4], grouping pathway models specific to SARS-CoV-2, other coronaviruses, as well as general cellular processes relevant for the virus-host interactions (e.g., ACE pathway, WP554[2]).

---

[2] https://www.wikipathways.org/index.php/Pathway:WP554

### Layout-aware systems biology formats

#### SBML

The Systems Biology Markup Language (SBML) [PMID:32628633] focuses on computational modelling of biological processes. SBML can store visual information about encoded elements and reactions using *render* [PMID:29605822] and *layout* [PMID:26528565] packages. An early version of SBML was adapted by CellDesigner [PMID:16082367] and introduced its specific encoding of layout and rendering information. When referring to SBML, we mean both layout+render SBML and CellDesigner SBML, indicating format-specific versions where necessary.

#### SBGN

Systems Biology Graphical Notation (SBGN) [PMID:19668183, doi:10.1016/B978-0-12-801238-3.11515-6] standardises visual encodings of molecular entities and their interactions. It facilitates a clear, common understanding of complex molecular mechanisms, allowing exchange and reuse of biological knowledge. SBGN is a graphical standard, and the corresponding SBGN Markup Language (SBGNML) [PMID:32568733; PMID:22581176] is a lightweight XML implementation of SBGN describing the layout of SBGN maps while preserving their underlying biological meaning.

SBGN consists of three complementary languages: Process Description [PMID:31199769] for a detailed description of molecular processes, Activity Flow [PMID:26528563] for encoding the high-level logic of molecular interactions, and Entity Relationship [PMID:26528562] for describing transitions between molecular states.

#### GPML

Graphical Pathway Markup Language (GPML) [PMID:25706687] offers a flexible, less-standardised representation of biological knowledge necessary to capture multiscale and, at times, ambiguous or uncertain biological knowledge. GPML is a structured XML format and can be systematically translated to other systems biology formats. Thus, GPML diagrams provide a qualitative link between the domain experts, pathway curators, and computational biologists.
