## Supplementary Material 3 for "COVID-19 Disease Map, a computational knowledge repository of SARS-CoV-2 virus-host interaction mechanisms"

### Description of bioinformatic resources


Tools for SBML curation and annotation (by Andreas Dräger and Alina Renz) 2

Creating interoperable content in SIF and SBML-qual format using CaSQ (by Anna Niarakis and Sylvain Soliman) 3

COVID-19 interaction dataset by the IMEx Consortium (by Pablo Porras) 4

COVID-19 interaction dataset by SIGNOR 2.0 (by Luana Licata) 5

Elsevier Pathway Collection (by Anastasia Nestorova) 5

OmniPath interactions database (by Denes Turei and Dexso Modos) 6

Pathway figure mining (by Alex Pico) 6

AILANI COVID-19 Text mining platform (by Dieter Maier and Angela Bauch) 7

OpenNLP-based text mining workflow (by Miguel Vázquez and Arnau Montagud) 8

The BioKB text mining platform (by Valentin Groues) 8

INDRA text mining platform (by Benjamin Gyori) 9

PathwayStudio text mining platform (by Anton Yuryev) 10

The COVIDminer project for visualisation of text mining networks (by Rupert Overall) 10

The BioKC platform for modelling-compliant curation (by Carlos Vega) 11

Leveraging COVID-19 Disease Maps for Counterfactual Inference of viral pathogenesis (by Jeremy Zucker) 12

Reactome Text mining and bioinformatic databases/Literature curation strategy (by Andrea Ribeiro and Robin Haw) 13

Atlas of Cancer signalling Network, ACSN (by Inna Kuperstein) 15

### Tools for SBML curation and annotation
#### (by Andreas Dräger and Alina Renz)

An adequate annotation of the model's components is required to enable the comparability and reuse of these models in SBML format. ModelPolisher [PMID:26882475] is a command-line based tool to annotate and autocomplete SBML models by accessing the manually curated BiGG Models knowledgebase [PMID:31696234]. It mainly relies on BiGG identifiers to recognize specific reactions and metabolites and extracts all available data about the identified component from the BiGG database [PMID: 31696234]. Its latest release added support for polishing models with identifier schemes beyond BiGG IDs. Moreover, ModelPolisher applies mappings from AnnotateDB [https://doi.org/10.5281/zenodo.3268443] for further annotations as well as appropriate Systems Biology Ontology (SBO) terms [PMID:22027554]. Besides the annotation of the model's components, ModelPolisher fixes apparent errors and performs inherent checks to provide a structurally correct SBML model. In the COVID-19 map project, ModelPolisher is used to enhance the model's annotations automatically.

The metabolic model test suite MEMOTE (memote.io) assesses the quality of genome-scale metabolic models in SBML format. Besides the structural validation of the file, MEMOTE benchmarks the model's annotations, biomass reaction, stoichiometry, and performs tests required for quality control [PMID:32123384]. Annotation tests check for the accordance with the minimum information required in the annotation of models (MIRIAM) guidelines [PMID:16333295] and for SBO terms [PMID:22027554]. ModelPolisher can increase the score of these tests drastically. MEMOTE makes it possible to track the development of the models from the project, and identifies further aspects for improvement.

In order to turn biochemical networks into dynamic models for simulation, kinetic equations need to be defined for every reaction. This process brings multiple problems with it. The first difficulty is choosing a specific rate law for a given reaction because of the large number of equations derived in more than 100 years of research in enzyme kinetics. Some equations come with thermodynamic pitfalls [PMID:20385728], while others require specific modulators to be present or a certain number of reaction participants. Second, the equations are often cumbersome to type and highly error-prone. Deriving the units for parameters and numerical values can be even more challenging. On a large scale, software support is indispensable for this purpose. SBMLsqueezer [PMID:26452770] is a versatile Java-based stand-alone tool. It comes with a graphical user interface, command-line interface, and an elaborate application programming interface that suggests and derives rate laws based on the structure and participants of biochemical reactions, including all units and parameters objects. In addition, retrieving experimentally obtained and manually curated rate equations with known parameter values from the reaction kinetics database SABIO-RK [PMID:29092055] has been an integral feature of CellDesigner [PMID:17822394] and is also implemented in SBMLsqueezer [PMID:26452770].

#### Creating interoperable content in SIF and SBML-qual format using CaSQ (by Anna Niarakis and Sylvain Soliman)

CaSQ [PMID:32403123] is a tool that converts static graphs built with CellDesigner (BIOSILICO, 1:159-162, 2003 [doi:10.1016/S1478-5382(03)02370-9]) to executable Boolean networks. Conversion rules and logical formulae are inferred according to the topology and the annotations of the starting molecular interaction maps that are in the form of Process Description (PD). The tool can process large and complex maps built with CellDesigner (either following SBGN standards or not) and produce Boolean models in a standard output format, SBML-qual, that can be further analysed using popular modelling tools. References, annotations, and layout of the CellDesigner molecular map are retained in the obtained model, facilitating interoperability and model reusability. Since version v0.7.8, the tool has been adapted to handle the notation schemes proposed for COVID-19 Map project and in the output files, the user can obtain two types of SIF files, one used for HiPathia (ref) modelling framework and one compatible with the CARNIVAL pipeline (PMID: 31728204). As the tool operates in two levels, graph conversion and rule inference, the outputs of the tool can be used in double sense: one for computational modelling as the SBML-qual files are fully executable, thus operational for performing in silico simulations, attractors search, and dynamical analysis and second, for obtaining through the produced SIF files, the structure of the regulatory graphs of the Boolean networks that consist of an AF-like graph derived from the Process Description (PD) molecular maps.

To sum up, we can obtain SBML-qual files for *in silico* simulations and analysis with CellCollective, GINsim or MaBoSS (see below), and Activity Flow simple interaction files (SIF) of the models, using two different naming schemes (regular names and raw aliases) facilitating interoperability and seamless downstream analysis of the COVID-19 Map content using both mechanistic and dynamic approaches.

#### COVID-19 interaction dataset by the IMEx Consortium (by Pablo Porras)

The IMEx Consortium is a group of databases such as IntAct [PMID:27115627], MINT [PMID: 22096227], MatrixDB [PMID: 30371822], UniProtKB [PMID: 30395287] and DIP [PMID: 11125102] that work together to represent experimentally detected physical molecular interactions from the literature using common formats and curation practices. IMEx expressive formats allow capturing interacting molecules and annotate them with binding kinetic parameters, sequence variation effects, cell line or tissue environments, or protein binding regions. The data is open and accessible. As a result of the COVID-19 pandemic, the IMEx Consortium curated Coronaviridae-related interaction data from reviewed manuscripts and preprints, resulting in a dataset of roughly 7300 interactions extracted from over 250 publications, including data from SARS-CoV-2, SARS, CoV, and other strains of Coronaviridae [doi:10.1101/2020.06.16.153817]. The dataset is updated with every release of IMEx data and can be accessed at the IntAct website (data:https://www.ebi.ac.uk/intact/query/annot:"dataset:coronavirus", data set description:https://www.ebi.ac.uk/intact/resources/datasets#coronavirus). Additionally, stable protein complexes from SARS-CoV-2 and SARS are being represented in the Complex Portal (https://www.ebi.ac.uk/complexportal) [PMID:30357405], a sister resource of IntAct that is used as reference for representation of multimolecular complexes by IMEx and other repositories. Both Reactome and WikiPathways currently cross-reference Complex Portal identifiers in their representation of SARS-CoV-2 and SARS related complexes.

#### COVID-19 interaction dataset by SIGNOR 2.0

#### (by Luana Licata)

SIGNOR 2.0, The SIGnaling Network Open Resource 2.0 (https://signor.uniroma2.it/) is an open-source database that manually annotates experimentally validated signalling data as binary causal interactions between biological entities (e.g. proteins, chemicals, phenotypes, complexes, drugs, etc.) [PMID: 31665520]. SIGNOR 2.0 annotates the effect (up/down-regulation) of a regulator entity on a target entity and the mechanism (e.g. phosphorylation, binding, transcriptional activation, etc.) behind the interaction.

Following the coronavirus pandemic, working in synergy with C19DMap Community, SIGNOR 2.0 started to organize in a structured format, information related to the host-virus interaction that causes COVID19 disease, in particular by annotating cellular pathways that are modulated during SARS-COV2 infection, such as apoptosis, inflammatory response and MAPK pathways.

The information was recovered from literature and involved causal interactions between SARS-CoV-2, SARS-COV-1, MERS proteins and the human host.

The resulting networks can provide a rational basis for the understanding of cellular perturbation during viral infection and to understand the mechanisms of action of SARS-CoV2. This information is available in a dedicated SIGNOR 2.0 web page (https://signor.uniroma2.it/covid/).

#### Elsevier Pathway Collection

#### (by Anastasia Nestorova)

The Elsevier Pathway Collection is the manually reconstructed dataset of annotated, interactive pathway diagrams (pathwaystudio.com). Statements about molecular interactions are extracted by Elsevier's text-mining technology [PMID: 15033866] into a knowledge graph supporting a range of relationship and element types [doi:10.1016/B978-0-12-817086-1.09989-9]. These interactions are curated into pathways having a rich list of annotations, including links to source literature, relevant database identifiers, and synonyms. This approach was adapted for extracting facts about viral proteins and viruses from the text, providing several thousands of COVID-19 related relationships from the literature. These interactions were filtered for experimental evidence and used for pathway reconstruction in the following categories: i) viral cycle and viral-human interactome for human SARS-CoV-2 and SARS-CoV-1; ii) cell-specific antiviral response; iii) sub-networks from bioinformatic analysis of transcriptomics and drug repurposing; and iv) complications associated with COVID-19.

#### OmniPath interactions database
#### (by Denes Turei and Dexso Modos)

OmniPath [PMID: 27898060] combines over 100 resources including pathways, protein interactions of inter- and intracellular signal transduction, as well as transcriptional and post-transcriptional regulation. This information is supported by rich annotations of protein roles including related diseases, function, or localisation. With OmniPath data, available via R/Bioconductor, Python, and Cytoscape [PMID: 31886476], SARS-CoV-2 infection mechanisms can be contextualised for a specific cell type by integrating network and omics data information [doi:10.1101/2020.08.03.221242]. Moreover, molecular details for relevant cell-cell interactions can be captured to trace the infection and immune reactions from primarily infected cells to other cell types.

#### Pathway figure mining
#### (by Alex Pico)

The vast majority of pathway knowledge is captured and communicated as image figures in the published scientific literature. Lacking a plain text representation or underlying data model, these figures are completely missed by text mining and informatic methods. With the goal of extracting pathway knowledge, the WikiPathways team performed optical character recognition on a collection of 64,643 pathway figures published over the past 25 years, followed by a customized text mining pipeline [doi:10.1101/2020.05.29.124503]. A total of 1.1 million human genes were found in these images (13,464 unique NCBI Gene identifiers); more content than is contained in text of the same papers and more than any pathway model database. Focusing on the COVID-19 Open Research Dataset (CORD-19) dataset, 221 pathway figures were mined and the results are presented as an online database that can be queried and browsed by gene symbol, disease associations, date and other metadata (https://gladstone-bioinformatics.shinyapps.io/shiny-covidpathways).

#### AILANI COVID-19 Text mining platform
#### (by Dieter Maier and Angela Bauch)

AILANI COVID-19 (https://ailani.ai) is an AI-based research assistant that combines semantic modelling, ontologies, linguistics and artificial intelligence (AI) algorithms. It uses natural language processing based literature mining and semantic mapping of structured databases and ontologies to generate knowledge graphs [DOI:10.1101/2020.04.17.044743]. We integrate and continuously mine the following resources: Medline, public PubMedCentral full-text articles, COVID-19/SARS-COV-2 bioRxiv/medRxiv articles, COVID-19/SARS-COV-2 Elsevier articles, ClinicalTrials.gov and relevant newsfeeds (e.g., WHO, CDC, NIH,…). In addition, structured data sources like Reactome or DrugBank, genome sequences, experimental and patent data are integrated.

We are applying AI on the knowledge graph (a semantic network incorporating ontology concepts, unique entity references to, e.g., genes/proteins/compounds, linked data from structured resources, and triples extracted from full-text documents)enables users to provide natural language queries and receive meaningful, high-quality answers. For example „What are COVID-19 risk factors?“ provides the answer „Hypertension, diabetes, COPD, cardiovascular disease, obesity, cerebrovascular disease, advanced age, immunosuppression, Lymphopenia and high CRP“ and complements classical extraction of knowledge from knowledge graphs by identifying yet unknown concepts

Classical keyword-based search with ontological drill-down to refine results allows interactive exploration. The "What are COVID-19 risk factors?" query retrieves almost 6000 individual hits which can be explored by association with specific diseases, drugs, pathways, etc.

For technical interoperability and batch processing, all information is also available by application programming interface (API). A REST Webservice is provided to execute AI, and keyword searches and specific endpoints allow to retrieve functions, pathways or drugs associated with submitted gene lists. Results are available in tab-delimited or JSON format (for technical documentation see online FAQ).

#### OpenNLP-based text mining workflow (by Miguel Vázquez and Arnau Montagud)

Text mining and natural language processing (NLP) were used to help curators and modellers complete their networks. The same pipeline was applied to two different corpora: One is the CORD-19, a collection of abstracts and full-texts compiled from COVID-19 related queries [PMID: 32510522], the other is the collection of MEDLINE abstracts associated to the genes in the PPI network from Gordon et al [PMID: 32353859] using the Entrez Gene reference into function (GeneRIF).

The text-mining pipeline consisted in identifying sentences using OpenNLP (https://opennlp.apache.org/) and mentions of genes using GNormPlus [DOI: dx.doi.org/10.1155/2015/918710]. GNormPlus tags genes with Entrez Gene identifiers, so no extra normalization was required. Sentences with mentions to at least two different genes were reported. Each of these sentences thus contains one or more pairs of genes that might have a protein-protein interaction (PPI) between them, for each of the corpora. Additionally, we built a tentative PPI network consisting of all the potential PPIs derived from each corpora, but restricted to only genes in the Gordon et al. publication. To help users, these 4 resources (2 for each corpora) contain the literal and normalized gene mentions, the text of the corresponding sentence, and the coordinates of those sentences in the original document.

#### The BioKB text mining platform (by Valentin Groues)

BioKB (https://biokb.lcsb.uni.lu) is a service available as a web application and as a SPARQL endpoint giving access to a knowledge base of biomedical facts extracted from the literature using a tailor-made text-mining pipeline. BioKB gives access to about 30 million facts (labeled relationships between entities) extracted from about 20 million publications. Additionally, named entities are systematically identified offering valuable insight into the frequency of co-occurrences. In the context of COVID-19, the text-mining pipeline was modified to better recognize the different terms related to the disease and to handle phenotype information using the Human Phenotypes Ontology. Moreover, the publications of the CORD-19 dataset were processed and a dedicated view was added to the web application: https://biokb.lcsb.uni.lu/topic/DOID:0080599

#### INDRA text mining platform (by Benjamin Gyori)

INDRA (Integrated Network and Dynamical Reasoning Assembler) [PMID:29175850] is an automated knowledge assembly system integrating information from published literature and biological pathway databases. Using multiple natural language processing systems ([PMID:30256986], Sparser[1], DRUM[2], [PMID:26357075]), INDRA processes relations extracted from natural language and pathway databases (e.g., TRRUST and Pathway Commons)[PMID: 29087512, 31647099] into a standardized representation called INDRA Statements. INDRA implements assembly algorithms to find exact and partial (e.g., hierarchical) redundancies between Statements from different sources, and calculates the overall confidence associated with each Statement using a probabilistic model. Each assembled INDRA Statement represents a unique biological mechanism linked to a collection of the underlying evidence text and PubMed ID or database entry references that support it. INDRA also implements model assembly modules to compile INDRA Statements into executable models (PySB, BNGL, Kappa, SBML), graphical standards (SBGN), Boolean/logical models, and multiple forms of causal interaction networks.

Using an automated content acquisition coupled to database storage on Amazon Web Services, INDRA processes all new literature appearing each day, and thus keeps up with COVID-19 literature in a timely manner. We systematically aligned the C19DMap with assembled INDRA Statements, both to enrich (i.e., find additional literature evidence for interactions already incorporated) and to extend (i.e., find relevant interactions that have not yet been incorporated) the C19DMap[3]. INDRA also supports natural language-based dialogue interaction with the COVID-19 literature (dialogue.bio), systematic causal path finding between factors (network.indra.bio), and reports on new knowledge collected from

---

[1] https://www.sift.net/publications/extending-biology-models-deep-nlp-over-scientific-articles
[2] https://www.aclweb.org/anthology/W15-3801.pdf
[3] https://github.com/indralab/covid-19/tree/master/covid_19/disease_maps

the COVID-19 literature each day[4]. INDRA also serves as the back-end to COVIDminer, as described in a section below.

#### PathwayStudio text mining platform (by Anton Yuryev)

Elsevier deep reading AI technology uses supervised machine learning to develop rules for automatic information extraction from text. Its entity name recognition layer uses various NER implementations, augmented by misspelled words recognition, part-of-speech tagging, and context sensitive name disambiguation.  Pattern matching algorithms are used to recognize genetic variation codes, protein modification sites and to handle combinatorial name explosion, e.g. in cell type and cell processes names.  Manually curated biomedical dictionaries and strict linguistic rules are used to reach a high level of accuracy rate for concept annotation (98%) and for relationship extraction (88%). These rates are similar to rates from manual annotation of the papers by experts [Elsevier R&D Solutions, 2018]. All synonyms and patterns are based on public database or ontology identifiers. Patterns for information extraction have single sentence scope and are written to match the text or semantic triples that are pre-extracted using a full-sentence syntactic parser.  ARelations extracted by pattern matching are named using a relation ontology based on the types of entities linked by relation and a verb used to specify a given relation.   Elsevier deep reading AI technology uses a variety of text and XML document formats as input, and it outputs extracted relations in RNEF XML format [doi: 10.1186/1471-2105-7-171] containing text snippet supporting relation to enable easy tracing to the original literature source. The Pathway Studio knowledge graph is updated  weekly. COVID-19 focused results of literature indexing can be accessed via the Elsevier Text Mining service [https://covid19.elsevier.com/].

#### The COVIDminer project  for visualisation of text mining networks (by Rupert Overall)

The COVIDminer project (https://rupertoverall.net/covidminer) maintains a biological interactome database generated from an automated reading of over 34000 COVID-19-

[4] https://emmaa.indra.bio/dashboard/covid19?tab=model

related manuscripts (plus more than 28000 abstracts where full text was not openly available). The text mining pipeline (based on the INDRA/REACH platform described above) is performed regularly to stay current in this extremely fast-moving field. Each extracted statement describes a directed interaction between two gene products, small molecules or biological processes. The causal network representing the COVIDminer database is browsable through an intuitive web interface. The ability to search the known literature for interactions (rather than just isolated keywords) and easily navigate the results, as well as rich links to supporting information and the original source manuscripts, make COVIDminer an invaluable tool for rapid literature curation. With well over 60000 manuscripts relating to COVID-19, curation is a task that can no longer be managed using traditional manual approaches alone. The global interaction network also enables the use of graph theoretical tools to extend maps or discover connections between existing pathways. COVIDminer is currently being used to help link the C19DM diagrams and build a more coherent picture of the disease as a whole.

#### The BioKC platform for modelling-compliant curation (by Carlos Vega)

BioKC is a web collaborative platform for the curation and annotation of systems biology facts. BioKC layout-less workflow provides quality control mechanisms for high quality curation of biological processes. Curators are provided with features to review the curation and annotation process, such as tasks, comments and an annotator agreement system to assess the task completion. Group managers can grant read, annotation, curation, or management permissions to control user roles. Elements composing facts can be assigned with supporting evidence from BioKB results or third party sources, keeping track of knowledge provenance. Moreover, BioKC allows for detailed annotation of the elements via uniform resource identifiers from 700 different namespaces available at identifiers.org. BioKC facts are encoded in SBML to ensure interoperability with other tools such as layout editors. Once a fact is ready to be released, it can be made public, entailing fact locking to prevent further changes, and the generation of a stable identifier which can be used as annotation in layout aware models. BioKC's role in the curation of COVID-19 disease maps is key to ensure high-quality model curation and help navigate the vast corpus of publications related to this novel disease.

### Leveraging COVID-19 Disease Maps for Counterfactual Inference of viral pathogenesis
(by Jeremy Zucker)

We are interested in leveraging domain knowledge contained in COVID-19 Disease Maps to enable three kinds of reasoning, collectively known as Pearl's Causal Hierarchy [Bareinboim et al 2020 https://causalai.net/r60.pdf]. The least expressive kind of reasoning is associational, which enables the ability to learn from data to answer questions such as "Given that we know a patient has COVID-19, would gaining information about her pre-existing conditions give us new information about her chances of survival?" The second level of the hierarchy is interventional, which enables the ability to learn from data to answer questions such as "Given that a patient has COVID-19, would treating her with Tocilizumab alter her chances of survival?" The third level of the hierarchy is counterfactual, which enables the ability to learn from data to answer questions such as "Given that I treated this COVID-19 patient with Tocilizumab and she lived, what are the chances that she would have died had I not treated her?" What enables this ability to reason at all three levels of the causal hierarchy is a formalism known as a Structural Causal Model (SCM) [Pearl 2000 http://bayes.cs.ucla.edu/BOOK-2K/]. An SCM represents domain knowledge in terms of causal diagrams, assumes a probability distribution over exogenous variables, and assigns deterministic functions to endogenous variables. Because the probability distribution over exogenous variables induces a probability distribution over endogenous variables, an SCM can answer associational level questions by using information about one endogenous variable to compute a posterior distribution on another endogenous variable. The action of replacing an endogenous variable's function with a fixed value enables the ability to learn from data to answer interventional level questions, and the process of using evidence to abduce a posterior distribution on exogenous variables followed by the action of fixing an endogenous variable enables the ability to answer counterfactual level questions. In order to leverage the COVID-19 Disease Map [PMID: 32371892] we are developing tools to translate a disease map into an SCM. We have recently developed an algorithm (BEL2SCM[5]) that automatically converts knowledge represented in the Biological Expression Language (BEL) [PMID: 24444544] to an SCM

[5] https://github.com/COVID-19-Causal-Reasoning/bel2scm

[Zucker et al 2020][6]. This same algorithm will work just as effectively with knowledge represented in the Systems Biology Graphical Notation Activity Flow (SBGN-AF) [PMID: 26528563]. We are also leveraging the ODE2SCM[7] methodology to generate an SCM for any ordinary or stochastic differential equation represented in the Systems Biology Markup Language (SBML) [PMID: 32845085]. Lastly, using the CASQ methodology [PMID: 32403123], it is possible to convert a disease map represented in Systems Biology Graphical Notation Process Description (SBGN-PD) [PMID: 31199769] to SBML Qualitative models (SBML-Qual) [PMID: 24321545], and we are now developing a tool to convert an SBML-Qual model to an SCM (QUAL2SCM).

#### Reactome Text mining and bioinformatic databases/Literature curation strategy
#### (by Andrea Ribeiro and Robin Haw)

With the rapid dissemination of COVID-19-related publications, we developed a triaging strategy to review and tag peer-reviewed publications to augment our traditional manual curation approach. In early 2020, literature metrics demonstrated that there were 30,311 articles and 5,638 preprints on SARS-Cov-2 (https://www.natureindex.com/news-blog/the-top-coronavirus-research-articles-by-metric; Chen et al., 2020). Our goal with triaging and tagging of SARS-CoV-2 publications was to provide a useful set of references for Reactome curators to assemble the specific molecules and molecular events into COVID-19-related pathways.

The literature triaging approach consisted of 4 main steps: i) Literature screening; ii) Literature selection; iii) Reference tagging; and iv) Reference database construction. For the literature screening approach, we mined different literature data sources. The main reference literature databases that were downloaded and parsed for step one (literature screening) were the CDC COVID-19 articles downloadable database (https://www.cdc.gov/library/researchguides/2019novelcoronavirus/researcharticles.html) and bioRxiv database (https://www.biorxiv.org/about-biorxiv), which also provided access to medRxiv references. Other collections of SARS-Cov-2 literature were also

[6] https://www.computer.org/digital-library/journals/bd/call-for-papers-special-issue-on-ai-for-covid-19
[7] https://github.com/robertness/ode2scm

manually screened, including:: the Zotero Library built and updated by members of COVID-19 Disease Map [PMID:32371892]; CORD-19 [PMID: 32510522]; LitCOVID [PMID:32157233]; and Johns Hopkins literature summary (https://ncrc.jhsph.edu/topics/); Cell Press Coronavirus Resource Hub (https://www.cell.com/COVID-19); Nature Coronavirus and COVID-19 updates (https://www.nature.com/collections/aijdgieecb); Science Latest Coronavirus research (https://www.sciencemag.org/collections/coronavirus?IntCmp=coronavirussiderail-128).

The initial process of selection of relevant references for Reactome SARS-CoV-2 map construction was based on title and abstract information. Within the first two months, rom 14221 references were screened in the CDC database.  700 (5%) articles were selected for further reviewing and tagging. Some of these initially selected articles were reje during the process of tagging as were not relevant in the context of our project. In the third step of our process around 290 (2%) references were tagged and integrated into the Reactome SARS-Cov-2 Library. The tagged references retrieved from the process of manual selection represented around 15% of the final Library. As Reactome is an evidence-based database built on reliable experimental published data, the process of SARS-CoV-2 reference selection was stringent.

SARS-CoV-2 Reactome tagging process was very detailed and specifically oriented for our purposes. Selected references were tagged regarding different features: (i) type of publication (i.e., article, review, pre-print, comment, opinion); (ii) Species (i.e., SARS-Cov-2, SARS-CoV-1; MERS, human ACE2); (iii) Entity (specific molecules studied); (iv) Methods (i.e., Cryo-EM, ELISA, binding assay, IC 50); (v) Cell line and/or Tissue (i.e., vero-E6, lung tissue); (vi) subcellular localization (i.e., plasma membrane, ER); Molecular event (i.e., virus cycle step, pathway, host response); and Phenotype (i.e., immune, coagulation, susceptibility). There was no limit to the number of tags and several included as many tags/words as necessary to provide a good overview of the reference to the Reactome curator.

Reactome SARS-Cov-2 Library was organized in a shared folder, where information was organized in a spreadsheet. Besides specific tags for each reference there are complementary information (DOI, Date of Publication, Title, PMID) and a link to the PDF file of the reference (stored in a shared folder). Importantly, Reactome members can edit and improve this Library as they used it (i.e., inclusion of tags).

Despite all the efforts of searching SARS-CoV-2 literature, the Reactome Library contained few references showing experimental data on molecular and therapeutic interactions. This is where Reactome's strategy of an individual map for SARS-CoV-1 map shows its value, supporting and guiding the construction of SARS-Cov-2 map. SARS-CoV-2 Reactome map is an ongoing task which will be built together with our understanding of the COVID-19 and in a fast pace as considered the scientific development and literature availability.

#### Atlas of Cancer signalling Network, ACSN (by Inna Kuperstein)

ACSN (https://acsn.curie.fr) is a web-based resource of multi-scale biological maps depicting molecular processes in cancer cell and tumor microenvironment. The core of the Atlas is a set of interconnected cancer-related signaling and metabolic network maps. Molecular mechanisms are depicted on the maps at the level of biochemical interactions, forming a large seamless network of molecular interactions leading the hallmarks of cancer. The Atlas is created using the systems biology standards and therefore is amenable for computational analysis. The maps of ACSN are organized in a hierarchical manner and decomposed into functional modules with meaningful network layout. Navigation of the ACSN is intuitive thanks to Google Maps-like features of NaviCell web platform. The processes represented on ACSN are often rather generic and also implicated in the COVID-19 pathophysiology, especially in stress response of host cell and in immune response. These molecular maps are updated and adjusted according to the literature about COVID-19 ("covidized") and thus included into the COVID-19 resource.
