## Supplementary Material 4 for "COVID-19 Disease Map, a computational knowledge repository of SARS-CoV-2 virus-host interaction mechanisms"

### Supplementary Material 4:
### Biocuration platforms and formats

#### Code and data availability

Code used for crosstalk analysis and the Cytoscape file with a selection of generated networks, are available in the public repository of the COVID-19 Disease Map project: https://git-r3lab.uni.lu/covid/models/-/tree/master/Resources/Crosstalks

#### Source data

##### Diagrams

Diagrams for the crosstalk analysis are from the COVID-19 Disease Map collection available at: https://covid.pages.uni.lu/map_contents. They were accessed as follows:

- Individual (MINERVA) diagrams via the MINERVA API: https://minerva.pages.uni.lu/doc/api/
- WikiPathway diagrams via the rWikipathways package: https://bioconductor.org/packages/release/bioc/html/rWikiPathways.html
- Reactome diagrams via the Reactome API: https://reactome.org/ContentService

##### Verified text mining interactions

Text mining interactions for enrichment of the existing diagrams were taken from the EMMAA Collection https://emmaa.indra.bio/dashboard/covid19?tab=model assembled using the INDRA platform (see Supplementary Material 2). Dataset timestamp: 2020-12-01-21-05-54.

Verified molecular interactions used for quality control of the text mining data were obtained from OmniPathDB (see Supplementary Material 2) using the OmnipathR package: https://bioconductor.org/packages/release/bioc/html/OmnipathR.html

We obtain verified text mining interactions by filtering the EMMAA dataset to interactions with “belief” of 0.8 or higher. Then, we retain only these interactions that match the direction and interacting molecules to the OmniPathDB dataset. We will call this filtered group of interactions “EMMAA-OP interactions”.

#### Crosstalk calculation

Crosstalks between source diagrams were calculated based on the HGNC identifiers of their elements. For simplification, all elements of the same diagram were considered to be interacting with each other. Three types of networks were constructed: existing crosstalks, new crosstalks and new regulators. To get a high-level perspective, diagrams can be grouped following the scheme in https://covid.pages.uni.lu/map_contents.

The networks were visualised using Cytoscape (https://cytoscape.org), with the Cytoscape Session File (.cys) available in the repository. The colour code is common for the networks: light green for pathway or group nodes, light blue for nodes having two neighbors, yellow for nodes having three or four neighbours, red for nodes with five or more neighbours. Pathway nodes have prefixes indicating their provenance. Pathway groups have no prefixes, as they combine diagrams across platforms.

##### Existing crosstalks

Existing crosstalks between diagrams, or groups of diagrams, were calculated by identifying shared HGNC identifiers. This way, we identify key elements that connect two or more different diagrams. The network was constructed as a graph composed of shared HGNC elements and nodes representing pathways or groups of pathways.

##### New crosstalks

New crosstalks between diagrams are new interactions, not present in current diagrams, that connect two or more different diagrams. To this end, we merge the EMMAA-OP interactions, (see above) with the network of existing crosstalks and keep only those new interactions that link at least two upstream and two downstream pathways, or pathway groups.

#### New upstream regulators

New upstream regulators between diagrams are new upstream elements, interacting with elements in current diagrams. To this end, we merge the EMMAA-OP interactions, (see above) with the network of existing crosstalks and keep only those new interactions with source elements not in existing diagrams, and target elements in at least one existing pathway or group.

### Crosstalk figures

Below are selected illustrations of crosstalk diagrams.

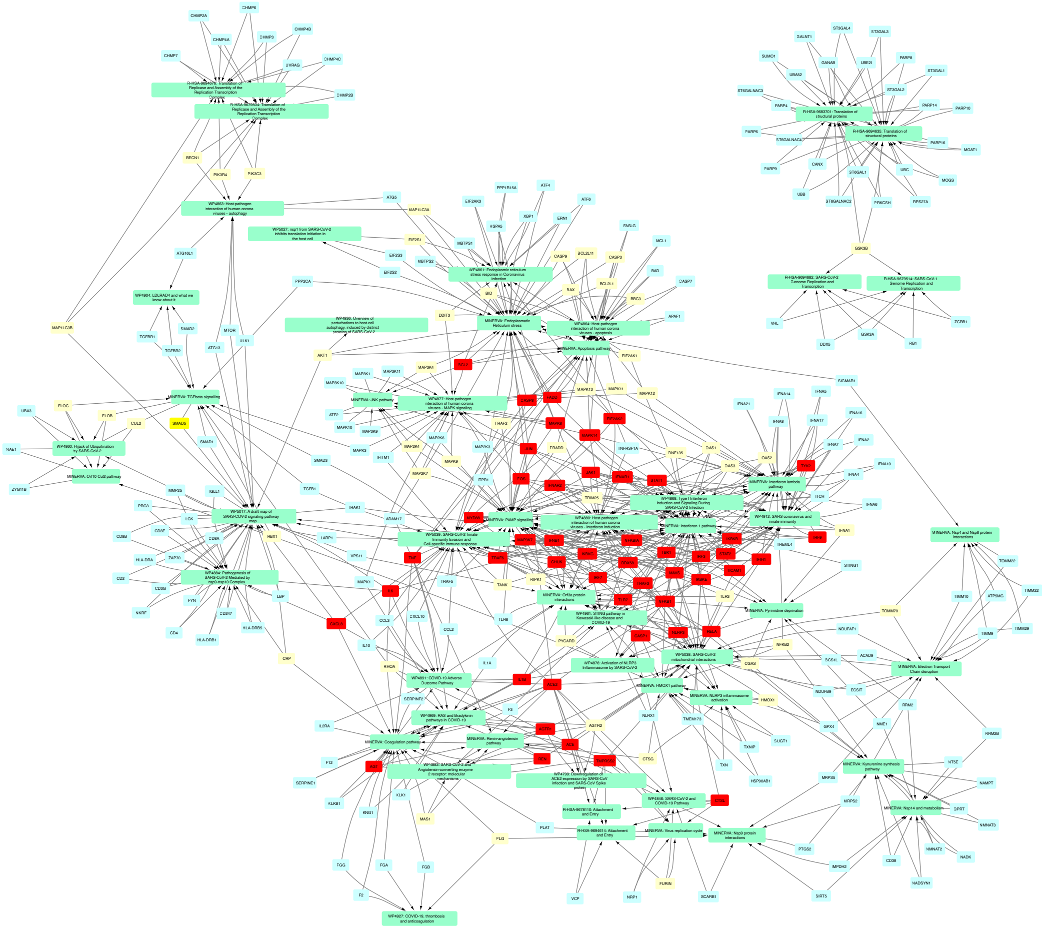

Fig S4.1 Existing pathway crosstalks

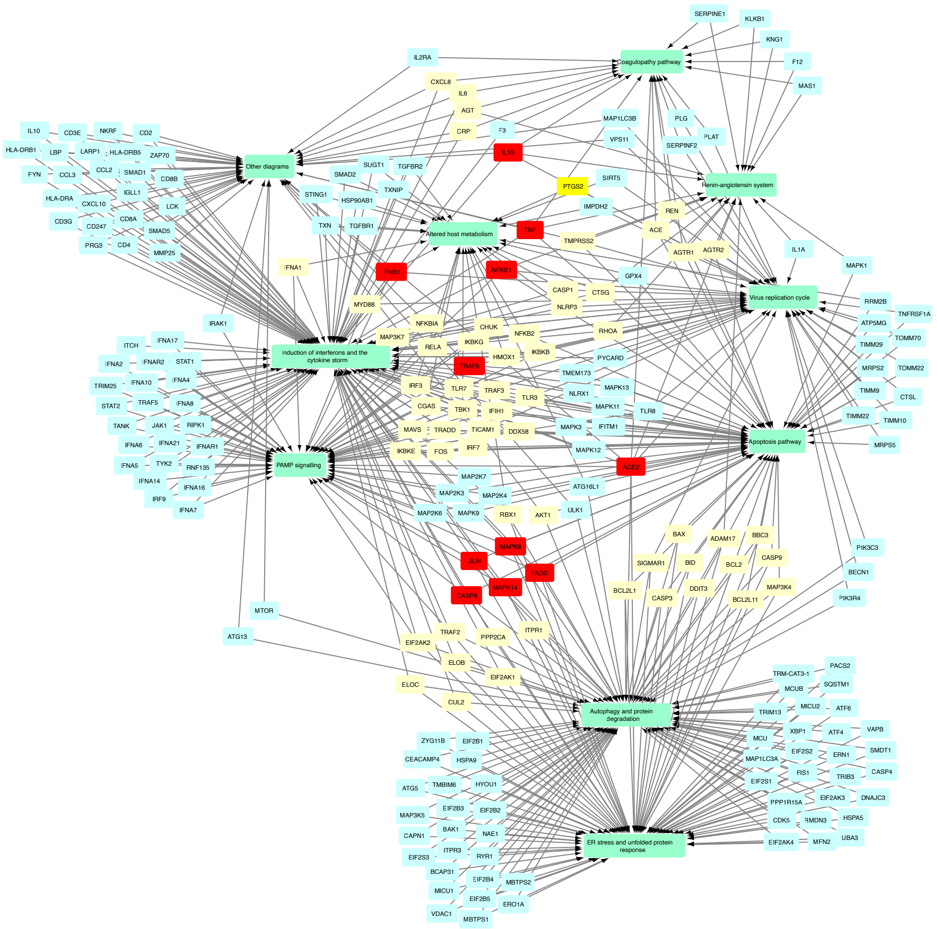


Fig S4.2 Existing group crosstalks

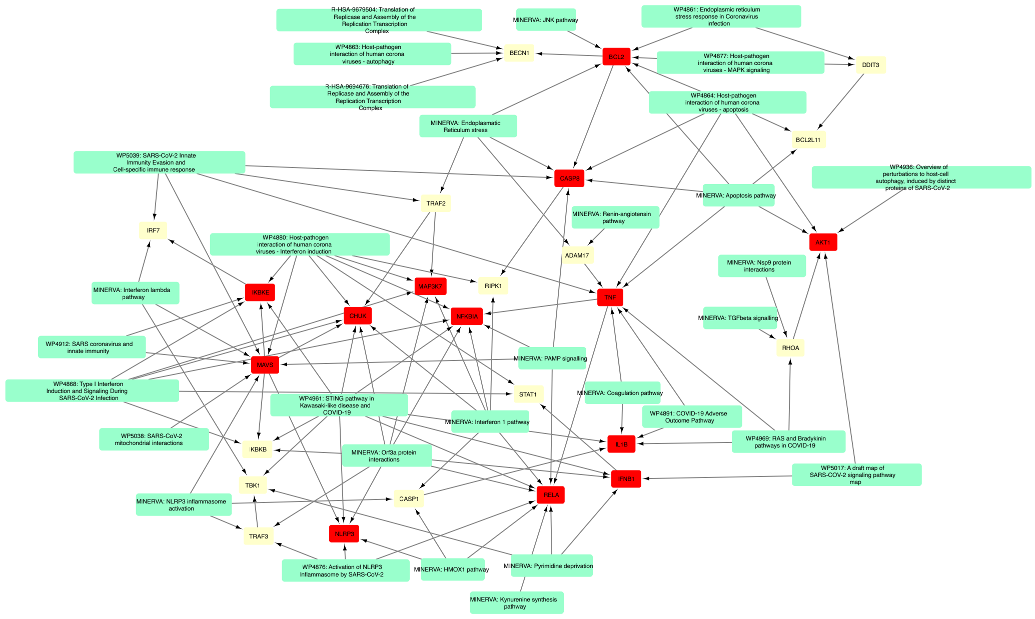


Fig S4.3 New pathway crosstalks

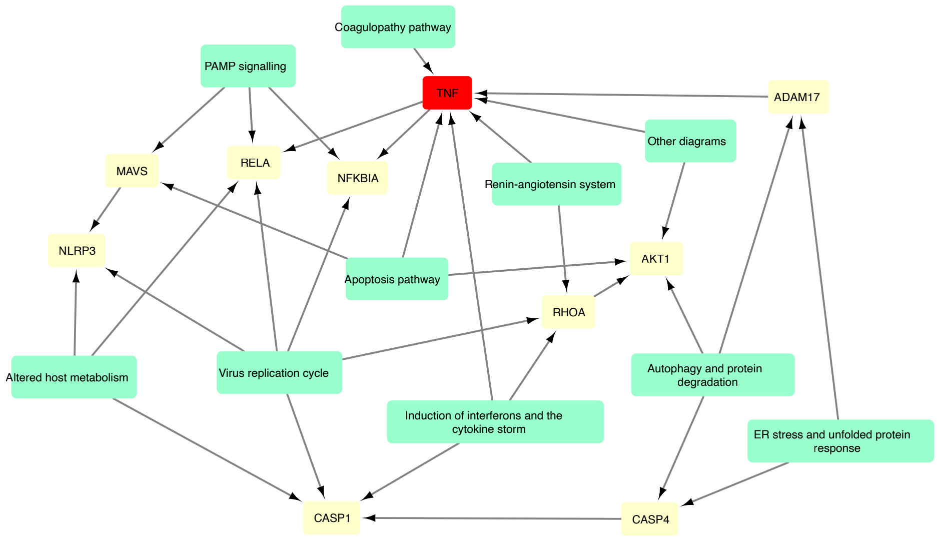


Fig S4.4 New group crosstalks

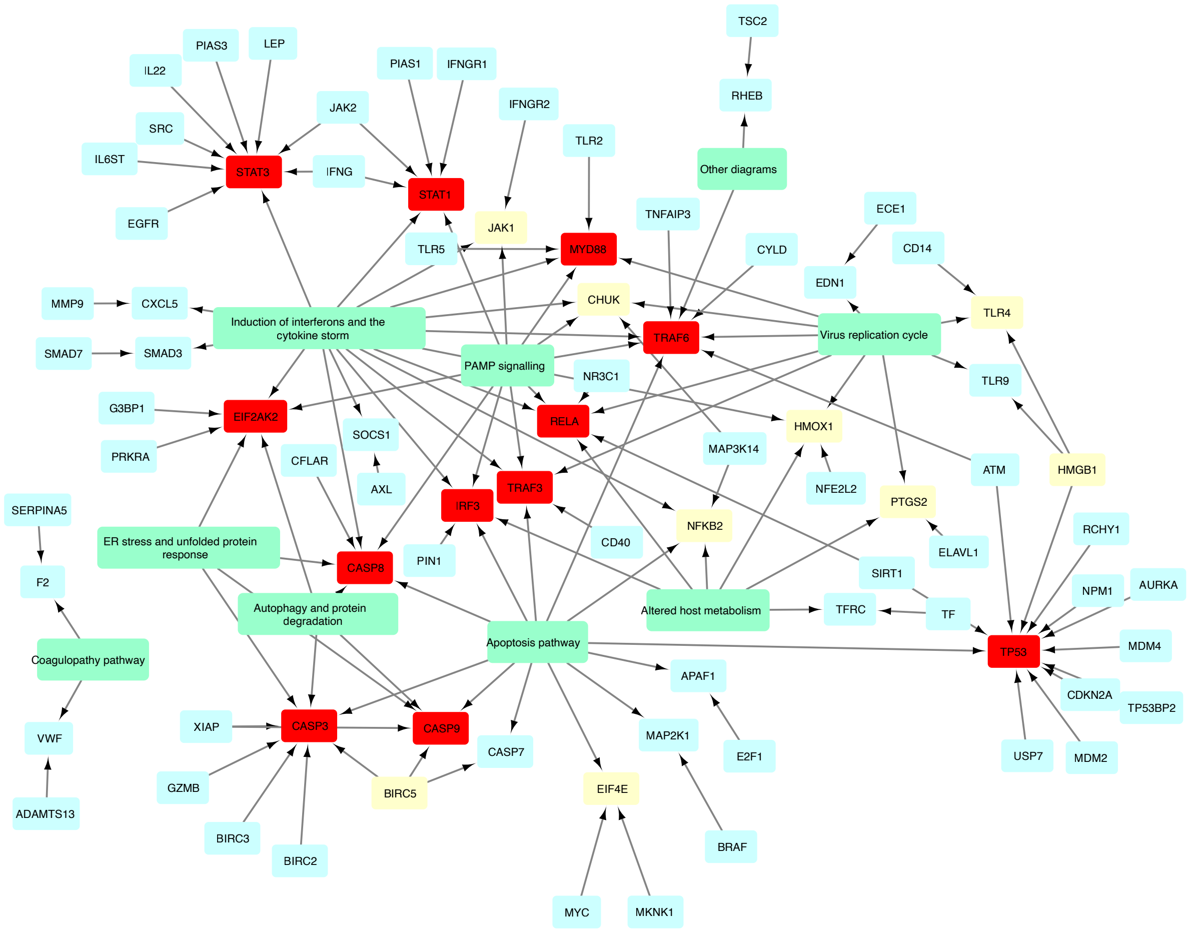


Fig S4.5 New group regulators
