## Supplementary Material 5 for "COVID-19 Disease Map, a computational knowledge repository of SARS-CoV-2 virus-host interaction mechanisms"

Hipathia results for the Apoptosis diagam of the COVID-19 Disease Map

| Subpathway_names | UP/DOWN | statistic | p.value | FDRp.value | Fold_Change | logFC |
|---|---|---|---|---|---|---|
| Apoptosis: Apoptosis | DOWN | -0,485661864 | 1 | 0,666474702 | 0,982756866 | -0,025093556 |
| Apoptosis: BAX | UP | 2,958122264 | 0,029154669 | 0,00185109 | 1,097857116 | 0,134690303 |
